# Fragment Screening and Structure-Guided Development of Heparanase Inhibitors Reveals Orthosteric and Allosteric Inhibition

**DOI:** 10.1101/2025.09.29.679132

**Authors:** Lani J. Davies, Cassidy Whitefield, Hyunjin Kim, Christoph Nitsche, Colin J. Jackson, Rebecca L. Frkic

## Abstract

Heparanase is the sole enzyme responsible for breaking down heparan sulfate within the extracellular matrix and its overexpression is linked to human diseases. Despite heparanase being a promising drug target, most efforts have focused on substrate mimetics, which have failed clinical trials, highlighting the need for new inhibitor scaffolds. Here, we employed fragment-based drug design to explore novel chemical space to develop small molecule inhibitors of heparanase. We used a crystallographic and computational approach to identify 31 fragments that bind heparanase; five of these inhibited heparanase in the micromolar range. One of these fragments underwent two cycles of fragment growing, which resulted in a compound with a seven-fold increased potency compared to the initial hit. The results from our fragment screen unveil untapped chemical space for heparanase inhibition, paving the way for the development of potent drug leads with the potential to transform the treatment of heparanase-related diseases.

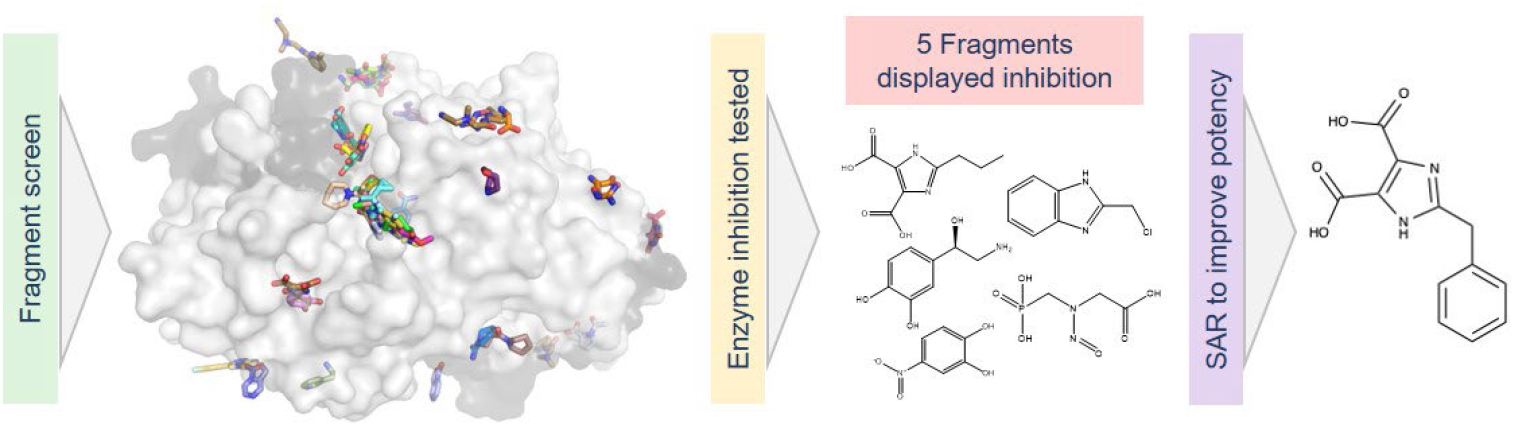

---

Heparanase is the only known mammalian enzyme that catalyses the hydrolysis of heparan sulfate in the extracellular matrix.^1, 2^ Under normal physiological conditions, the expression of heparanase is strictly controlled to prevent unintended tissue damage.^2^ However, overexpression of heparanase in disease accelerates the degradation of heparan sulfate, which can have detrimental effects.^3, 4^ For example, heparanase is upregulated in a number of inflammatory diseases including Crohn’s disease, ulcerative colitis, rheumatoid arthritis, chronic pancreatitis, diabetes, and diabetic retinopathy.^5^ Importantly, heparanase is overexpressed in every human cancer that has been studied to date, where it is associated with an increase in metastasis, tumour size and tumour angiogenesis.^1, 6, 7^

The overexpression of heparanase in a number of diseases presents an opportunity to target heparanase with inhibitors to reduce the severity of disease.^8^ However, identifying effective drugs for heparanase inhibition remains a challenge and there are no highly specific, clinically approved heparanase inhibitors.^9^ As the natural substrate for heparanase is heparan sulfate, a complex and highly charged polysaccharide, the common choice of starting molecules for drug development are often oligosaccharides that mimic this substrate, such as pixatimod (PG545), muparfostat (PI-88), roneparstat (ST0001) and necuparanib (M402).^10-13^ However, the clinical trials of each of these mimetics were unsuccessful due to side effects or inefficiency, and none have been successful in gaining approval as heparanase inhibitors.^11, 12, 14-20^ In contrast to heparan sulfate mimetics, small molecules have a greater chemical diversity and can be more easily optimised through structure-activity relationship studies to enhance pharmacokinetic properties and oral absorption, while also minimising off-target effects through optimising specificity.^21^

Although several small molecule active site inhibitors of heparanase have been identified, only a few have advanced to clinical trials, with none progressing beyond this stage.^5, 22^ Among the early candidates were OGT-2115 and OGT-2492 (IC_50_ of 3.0 µM and 0.4 µM, respectively).^13^ Another compound, suramin, a synthetic polysulfonated naphthylurea originally developed as an antitrypanosomal drug, exhibited non-competitive inhibition of heparanase with an IC_50_ of 46 µM.^23^ Suramin progressed to clinical trials as a heparanase inhibitor, however this was halted due to severe side effects.^23^ RK-682, a 3-alkanoyl-5-hydroxymethyl tetronic acid derivative, also showed inhibitory activity with an IC_50_ of 17 µM, but did not proceed to clinical development.^24^ Irreversible heparanase inhibitors inspired by the natural glycosidase inhibitor cyclophellitol have also been shown to reduce cancer aggression with IC_50_ values determined by colorimetric assay to be as low as 0.46 µM.^25, 26^ Similarly, iminosugar inhibitors derived from siastatin B, also a glycosidase inhibitor, are potent heparanase active site inhibitors with a reported IC_50_ value of 27 µM.^27^ Most recently, a known antagonist of the G-protein coupled receptor LPA-5, TC LPA5 4, has displayed a reported heparanase IC_50_ of 10 µM.^22, 28^

The chemical diversity of known heparanase inhibitors is relatively limited, which impedes the efficient development of effective small molecule inhibitors through analogue-based approaches.^21^ Fragment-based drug discovery (FBDD) offers a promising strategy to maximise the coverage of chemical space during screening campaigns. FBDD has a high success rate in generating a series of lead molecules for drug development when compared to traditional high-throughput screening.^29^

In this work, we applied the principles of FBDD through a combination of structural characterisation using X-ray crystallography, as well as computational docking and inhibition assays to identify 31 novel ligands of heparanase. We subsequently increased the potency of one of these fragments using fragment growing strategies. This work enhances our understanding of the chemical diversity of heparanase inhibitors and identifies key chemical motifs that are essential for effective heparanase binding.

We utilised our engineered human heparanase variant^30^, which shows the same structure, activity, and dynamics as the wildtype but enables recombinant expression in *E. coli*, with a commercial library of 96 fragments to discover initial chemical moieties that bind heparanase. Each fragment was soaked into apo-heparanase crystals, and we obtained datasets of acceptable resolution (<2.6 Å) for 92 of the 96 fragments. To streamline fragment identification in our datasets, and to identify weak binding events, we utilised the Pan-Dataset Density Analysis (PanDDA) software.^31^ The inclusion of 32 apo datasets (Supplementary Table 1) in addition to our 196 putative fragment-bound datasets enabled the PanDDA software to calculate and subtract background solvent and noise levels to detect ligand binding in a high-throughput manner. We discovered 27 fragments had bound to heparanase, with several fragments occupying two or more binding sites (Figure 1 and Supplementary Table 2).

**Figure 1:**
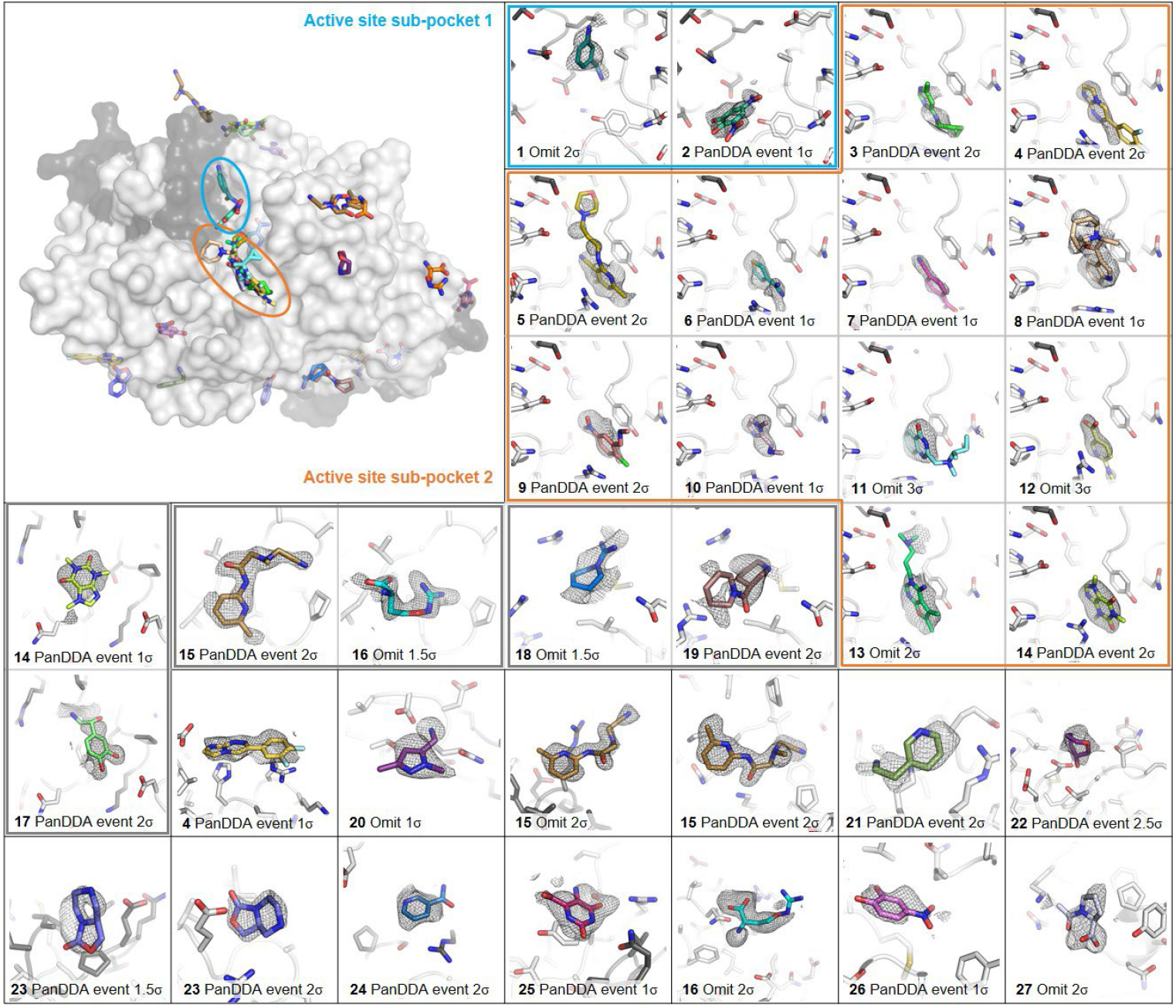
Fragments bound to heparanase from the crystallographic fragment screen. (top left) heparanase shown as surface with large subunit coloured in light grey and small subunit coloured darker grey. Bound fragments are shown as coloured sticks. (all other panels) Fragments bound in the active site sub-pocket 1 (contained in blue outline), active site sub-pocket 2 (orange outline), and remote fragments. Grey boxes are around remote fragments that share the same binding site. PanDDA event maps and mF_o_-DF_c_ omit maps are shown in grey mesh, with contour levels indicated for each panel. Fragments are shown in coloured sticks, and contributing side chains of heparanase are shown in grey sticks.

Of the 27 bound fragments, 14 (compounds **1** – **14**) are located within the active site in two primary sub-pockets (Supplementary Table 3). The smaller “sub-pocket 1” contained two fragments (**1** and **2**). **1** is located deep in the binding pocket comprised of the interface between the small and large subunits of heparanase and interacts with the enzyme through predominantly hydrophobic interactions with Ala388 of the large subunit and Asn64 and Leu65 of the small subunit. Fragment **2**, however, forms more specific interactions with side chains lining the binding site, including hydrogen bonds with Tyr391 and Gln383, as well as water-mediated bonds with the backbone of the protein. Despite occupying the same pocket, there is no spatial overlap between the binding modes of these two fragments, suggesting a potential avenue for fragment linking.

“Sub-pocket 2” is a larger hydrophobic pocket situated amongst the catalytic residues Glu225 and Glu343, and contained the greatest abundance of fragments bound. The 12 (compounds **3** – **14**) fragments bound in sub-pocket 2 are generally larger than the other fragment hits and share common aromatic motifs that align well in the same plane and exhibit π-π stacking with Tyr348, and some short substituents that extend to interact with the catalytic residue Glu225.

The remaining 19 fragment binding events (compounds **14** – **27**) were dispersed amongst remote sites. While most remote binding events were isolated to one fragment at the site, three pairs of fragments occupied common binding sites (Figure 1, grey boxes); **15** and **16** both occupied a shallow surface pocket between Pro353 and Thr306, **14** and **17**, whose similar size allowed them to fit in a shallow pocket between Arg70 and Pro78 of the small subunit and Asn397 of the large subunit, and **18** and **19**, which showed almost perfect overlap of their six-membered rings in a binding cleft between Gln324 and Arg374, providing an avenue for fragment merging (Supplementary Figure 1). The remote binding fragments displayed greater diversity in ligand chemistry compared to fragments that bound to the active site and interacted with heparanase through a combination of hydrophobic and polar interactions.

To sample an even larger chemical space to discover new molecules to bind heparanase, computational fragment screening was conducted in parallel with the crystallographic screen. Schrödinger’s Maestro^32^ suite of programs were used to identify potential binding sites on heparanase based on crystal structures. SiteMap^33^ analysis identified the active site plus five distinct potential allosteric sites suitable for further docking studies. Interestingly, the remote sites identified by SiteMap did not overlay well with the remote binding sites uncovered in the crystallographic fragment screen (Supplementary Figure 2). Nevertheless, the “Fragmenta” (http://www.fragmenta.com/) library comprising 2099 fragments was systematically docked into each of the computationally-identified binding sites using increasingly stringent docking methods. A final set of 18 commercially available fragments across the six identified sites were pursued based on their docking G-score and commercial availability (Supplementary Table 4, Supplementary Figure 3).

In order to experimentally validate the binding poses of the computationally discovered fragments, we used X-ray crystallography (Supplementary Table 5). We employed the same soaking method as the commercial crystallographic screen to obtain suitable datasets for 17 of the purchased fragments. Among the datasets assessed, there was density identified using PanDDA for four fragments (compounds **28** – **31**), with one fragment, **28**, bound in two sites (Figure 2). Three unique fragments were bound in the active site, two in active site sub-pocket 1 (**28** and **29**) and two in sub-pocket 2 (**28** and **30**), while the fourth fragment binds in a small pocket adjacent to the active site (**31**).

**Figure 2:**
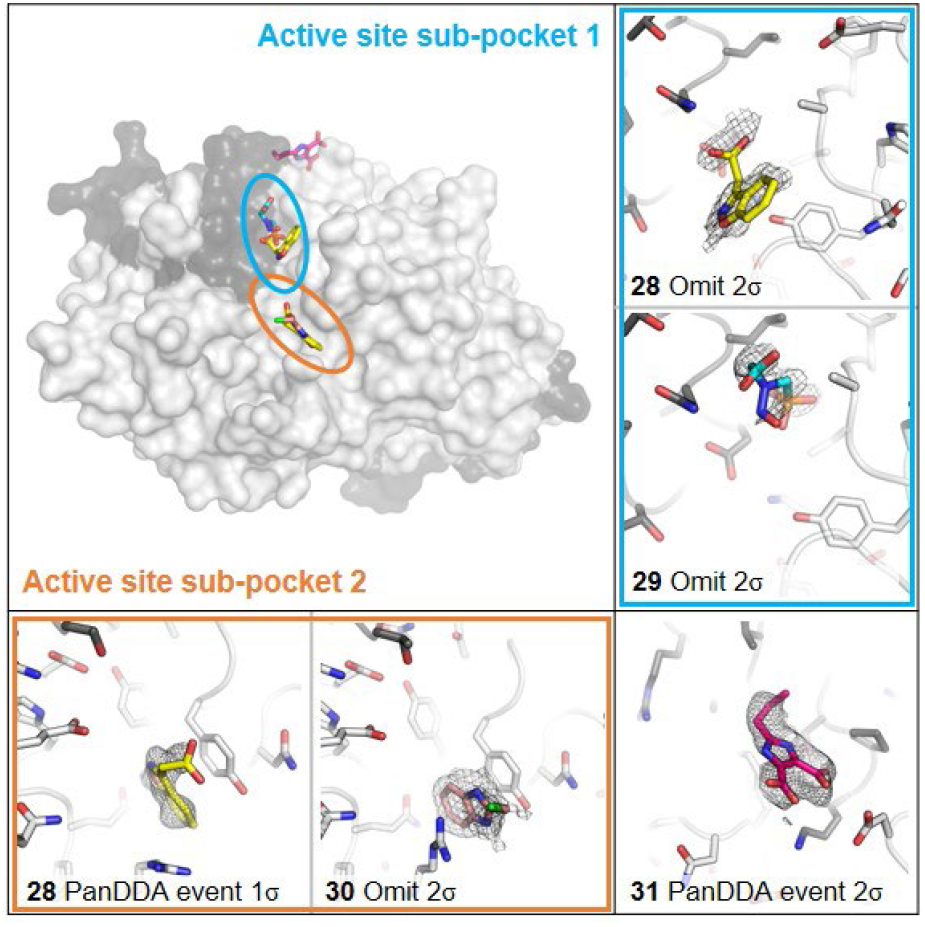
Fragments bound to heparanase from the computational fragment screen. (Top left) heparanase shown as surface with large subunit coloured in light grey and small subunit coloured darker grey. Bound fragments are shown as coloured sticks. (all other panels) Fragments bound in the active site sub-pocket 1 (contained in blue outline), active site sub-pocket 2 (orange outline), and allosterically. PanDDA event maps and mF_o_-DF_c_ omit maps are shown in grey mesh, with contour levels indicated for each panel. Fragments are shown in coloured sticks, and contributing side chains of heparanase are shown in grey sticks.

We observed little agreement between the computationally predicted binding sites and the crystallographically determined binding modes, with **29** being the only fragment to bind at its predicted site. **28** and **30** were predicted to bind remotely but bound in the active site, and **31** was predicted to bind in the active site but bound remotely in our crystal structure. This highlights the promiscuity of fragment binding, and that additional factors such as protein flexibility and solvent effects can influence their final binding mode. Additionally, our crystal structures highlight the necessity of experimental validation in fragment-based drug discovery, as computational predictions alone are not reliable in determining actual binding interactions.

Interestingly, the remote binding site of fragment **31** was the same as that of **14** and **17** from the crystallographic fragment screen, with their analogous moieties overlapping in their pose (Supplementary Figure 4). Their aromatic rings were positioned to form *π-π* stacking with Phe398 and **17** and **31** formed a hydrogen bond with Ser77 of the small subunit. Large differences between the chemical structures of the three fragments account for very little change in their binding modes, due to relatively non-specific interactions occurring with the protein at this site.

To assess the capacity for the fragments to inhibit heparanase, their effect on heparanase activity was measured using a colorimetric assay developed by Hammond et al.^30, 34^ This assay monitors the heparanase-mediated cleavage of fondaparinux where fragments were tested at concentrations up to 5 mM. For those that showed inhibitory activity, the maximum inhibition was calculated based on the reduction in heparanase activity, and IC_50_ values were determined as the concentration required to achieve 50% of this maximum inhibition (potency). 18 fragments that showed binding in the crystal structures were selected for activity testing based on commercial availability, and five of these displayed heparanase inhibition when tested up to 5 mM (Table 1). Potencies ranged from 142 – 970 µM IC_50_, with fragment **17** identified in the crystallographic screen showing the lowest IC_50_.

**Table 1.**
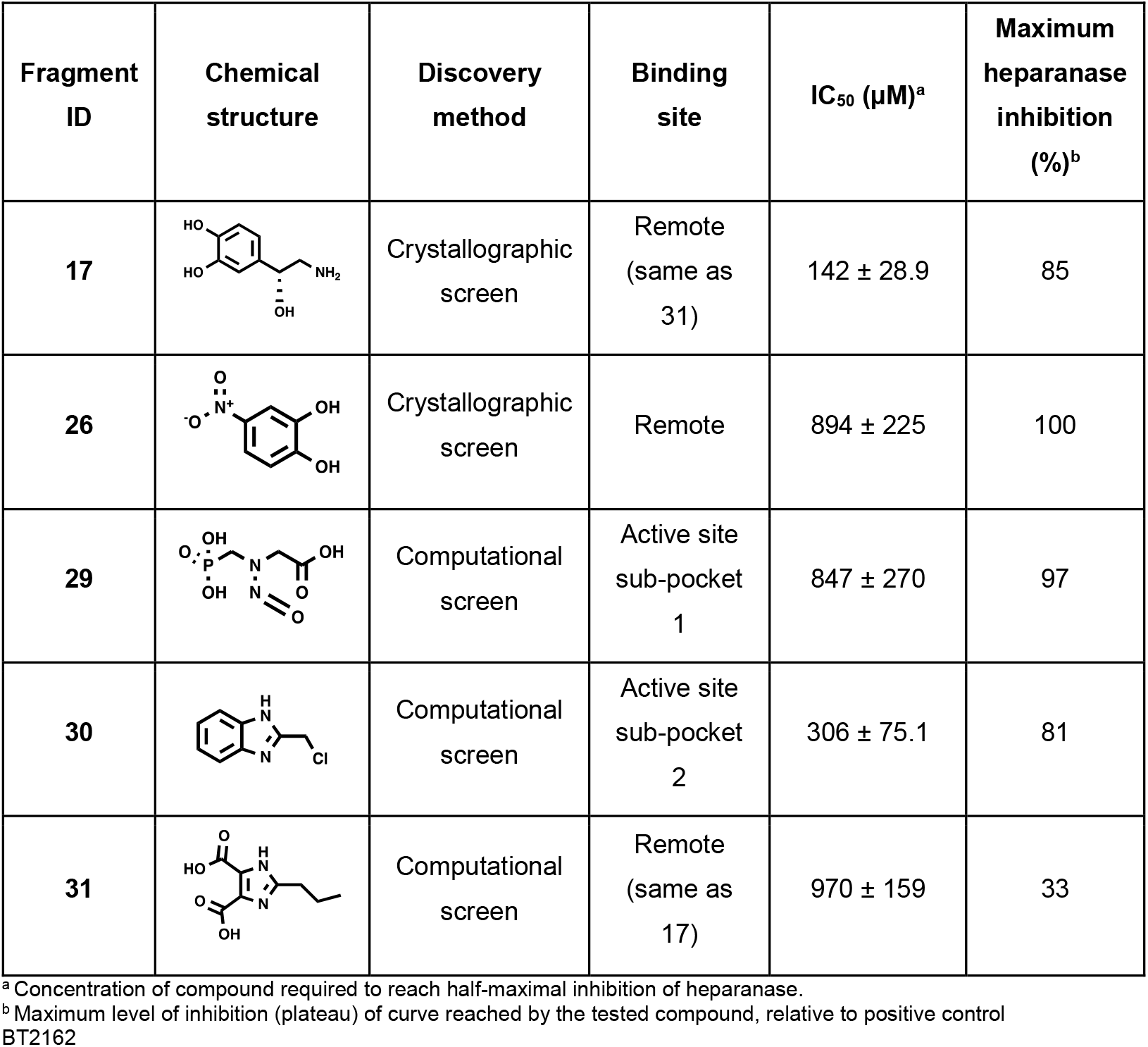
Inhibitory activity displayed by fragments binding to heparanase.

| Fragment ID | Chemical structure | Discovery method | Binding site | IC<sub>50</sub> (µM)<sup>a</sup> | Maximum heparanase inhibition (%)<sup>b</sup> |
| --- | --- | --- | --- | --- | --- |
| **17** |  | Crystallographic screen | Remote (same as 31) | 142 ± 28.9 | 85 |
| **26** |  | Crystallographic screen | Remote | 894 ± 225 | 100 |
| **29** |  | Computational screen | Active site sub-pocket 1 | 847 ± 270 | 97 |
| **30** |  | Computational screen | Active site sub-pocket 2 | 306 ± 75.1 | 81 |
| **31** |  | Computational screen | Remote (same as 17) | 970 ± 159 | 33 |

<sup>a</sup> Concentration of compound required to reach half-maximal inhibition of heparanase.
<sup>b</sup> Maximum level of inhibition (plateau) of curve reached by the tested compound, relative to positive control BT2162

Two out of the five active fragments were bound in the active site. These fragments, **29** and **30**, were identified in our computational screen and were bound in active site sub-pocket 1 and 2, respectively, and showed modest inhibition of heparanase with IC_50_ values of 847 and 306 µM, respectively. **29** occupied the active site sub-pocket 1 and inhibited heparanase activity by 97% but with a lower potency of 847 µM. **1** and **24**, which were identified in the crystallographic screen but showed no inhibitory activity, also occupied this site and are chemically similar to each other but distinctly different from **29**. This suggests greater binding promiscuity in the pocket than initially indicated by the crystal screen, and the scaffold of **29** is a better starting point for inhibitor design. However, **29** has predicted toxicity^35-37^ and so we did not pursue this fragment further. The most potent of the computationally discovered fragments was **30**, which displayed an IC_50_ of 306 µM. This fragment occupied the active site sub-pocket 2 and inhibited heparanase activity by 81%. It bound in a similar way to the active site sub-pocket 2 fragments from the crystallographic fragment screen, where it forms a π-π stacking interaction with Tyr348. It is unclear why only fragment **30**, among those bound in active site sub-pocket 2, exhibited activity against heparanase.

Three of the five fragments showing inhibitory activity against heparanase were bound in two remote sites in our crystal structures, indicating potential allosteric mechanisms of inhibition. Fragment **17** displayed the lowest IC_50_ of the tested fragments and was bound on a shallow remote surface pocket, stabilised by hydrogen bonds with Ser77 and the backbone carbonyl of Ile73, as well as a π-π stacking interaction with Phe398 (Figure 1). This fragment displayed 85% maximum inhibition of heparanase activity, with an IC_50_ of 142 µM. Interestingly, **14** and **31** co-localised with **17** at this binding site (Supplementary Figure 4) where **31** but not **14** also showed inhibitory activity. **14** did not form specific hydrogen bonding interactions with heparanase, suggesting subtle differences in specific interactions with heparanase can enable binding but not inhibition by two similarly sized but chemically diverse fragments. **31** was the only computationally identified fragment that did not bind in the active site in our crystal structures. In fact, the effect of **31** on heparanase activity was the lowest of the inhibitors, with a maximum enzyme inhibition of 33% and an IC_50_ of 970 µM. Despite their differing chemical structures, both **17** and **31** align well in their binding modes, positioning key functional groups similarly within the pocket (Supplementary Figure 4). The activity displayed by **17** and **31**, despite their chemical diversity, offers a strong foundation for developing inhibitors probing the same binding site with potential for allosteric inhibition.

**26** was the only other active fragment identified from our crystallographic screen, and was also shown to not bind to the active site. **26** bound in a deeper pocket than **17**, forming a hydrogen bond with the backbone carbonyl of Ile237. The binding mode of **26** caused a large displacement of the side chain of Met278 compared to all other structures (Supplementary Figure 5); this appeared to be an isolated conformational change that did not propagate through nearby residues. This fragment displayed higher activity against heparanase than **17**, with a maximum inhibition of 100% but with lower potency (IC_50_ = 894 µM). The binding site of **26** is promising for the development of an allosteric inhibitor, as it was the only one not bound to a shallow surface pocket. Instead, **26** occupies a cleft generated by a displacement in Met278, offering potential for fragment elaboration to enhance binding site specificity and increase potency.

While the hit rates for binding were similar between the crystallographic (29%) and computational (22%) screens, a much higher proportion of the tested fragments that displayed inhibitory activity were computationally derived (75% vs. 14%). This suggests that the computational screen, tailored to heparanase’s binding sites, identified more active and specific hits, despite minimal overlap between predicted binding poses and crystallographically captured binding poses. This discrepancy highlights the limitations of docking in accurately predicting binding modes while still enriching for functional inhibitors, suggesting that computational screening is valuable for identifying potent fragments but should be complemented with structural validation.

In an effort to increase the potency of the fragments for inhibiting heparanase, a Structure Activity Relationship (SAR) by catalogue study was carried out to generate grown analogues of the active fragments. The four fragments identified as inhibitors (excluding **29** due to predicted toxicity) underwent computational SAR studies. Initially, the original fragment structures were input into Scifinder to explore commercially available molecules sharing the same scaffold. A catalogue of grown fragments was computationally docked to the binding site of their parent hit. In most cases, the grown fragment either could not be docked to the binding site of the original fragment due to space constraints within the pocket, or returned poor docking scores indicative of unlikely binding.

We purchased five grown fragments and tested their activity against heparanase. Of these, only one showed activity against heparanase – **32**, a grown analogue of fragment **31** (Figure 3A). This fragment showed a three-fold improvement in potency compared to the original **31** fragment, exhibiting an IC_50_ of 319 µM (Figure 3B). In addition to higher potency, the grown fragment displayed higher inhibitory activity against heparanase, with a maximum inhibition of 74%.

**Figure 3:**
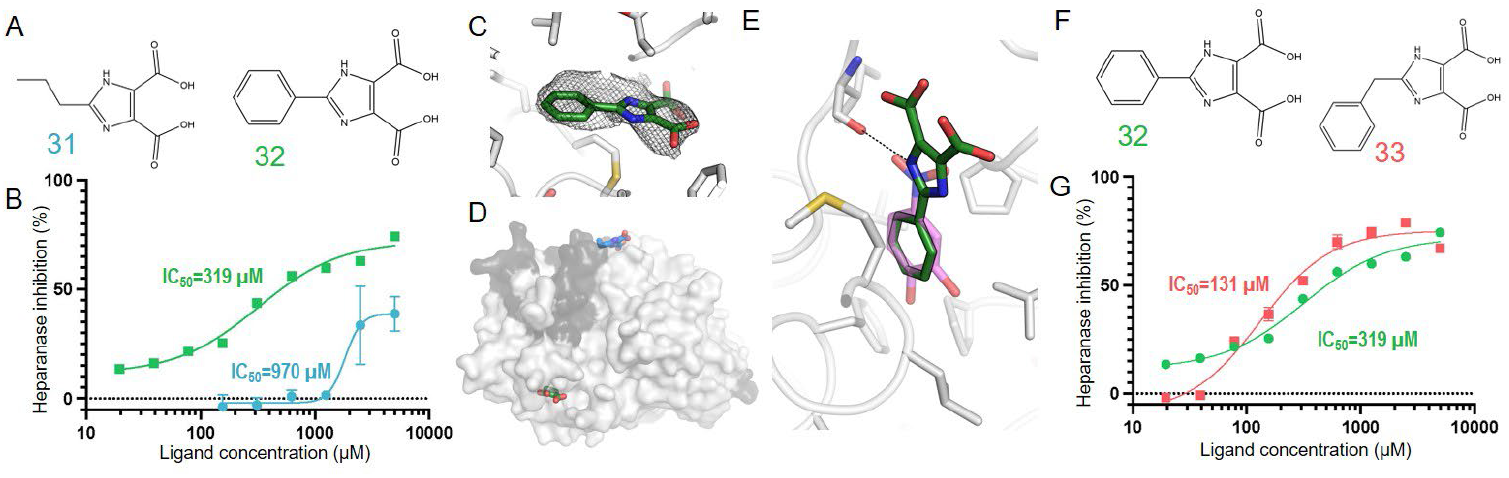
Activity and structural mechanism of grown fragment 31. (A) Fragment **31** underwent SAR by catalogue to generate a grown fragment **32**. (B) Inhibition curves of the original fragment, **31** (blue) compared to the grown fragment, **32** (green). (C) PanDDA event map for **32** ligand binding shown in mesh, contoured to 2σ. (D) Differences in binding sites of **31** (blue) and **32** (green). Heparanase is shown as surface, the large heparanase subunit coloured light grey, and the small subunit coloured dark grey. (E) Comparison of binding modes of **32** (green sticks) and **26** (translucent pink sticks), which both cause a conformational displacement in Met278. (F) Secondary SAR by catalogue produced **33**. (G) Inhibition curves of **32** (green), compared to second generation grown fragment **33** (pink).

To determine the mechanism behind the increased activity of **32**, we obtained an X-ray crystal structure of **32** bound to heparanase (Figure 3C, Supplementary Table 6). **32** bound in a different allosteric site to the original **31** fragment (Figure 3D), and instead bound to the same cleft as **26**, while also causing a displacement in Met278. The common phenyl ring of the two molecules overlap well in the cleft and similarly to **26, 32** forms a hydrogen bond with the backbone carbonyl of Ile237 (Figure 3E). The potency of **32** was higher than **26**, which exhibited an IC_50_ of 894 µM. This could be attributed to the absence of polar additions on the phenyl group of **32**, which enables favourable interactions with surrounding hydrophobic residues, including Pro226, Gly239, Leu242, Pro266, Val268, Leu279, and Phe282. Conversely, the addition of two hydroxyl groups on the phenyl ring of **26** likely creates a comparatively unfavourable electrostatic binding environment, reducing the potency of **26** in comparison to **32**.

Encouraged by the improvement in potency observed in a single round of SAR, we sought to undergo a second iteration of fragment growing to determine whether the IC_50_ could be further reduced. We used a similar approach as the first round to generate a library of grown fragments that were docked to the same binding site as **32**. We purchased a selection of six grown fragments and tested their activity against heparanase which revealed that **33**, a grown fragment with an extended linker to the phenyl group (Figure 3F), showed an improved potency of 131 µM IC_50_ and maximum inhibition of 80% (Figure 3G). Despite the increased potency, we were unable to obtain a crystal structure of this grown fragment in complex with heparanase. This is likely due to the addition of a carbon atom in the linker connecting the phenyl group to the imidazole ring, increasing the hydrophobicity of the compound and subsequently reducing its solubility and ability to co-crystallise. Nonetheless, the demonstrated improvement in potency of the computationally discovered fragment **31** by more than seven-fold (970→131 µM) using fragment growing strategies is a viable proof of concept to establish fragment-based drug discovery as a promising avenue for the development of heparanase inhibitors.

Despite our drug discovery efforts being carried out using an engineered heparanase variant, our results are likely to be of relevance to the wild-type human heparanase, as superimposition of the fragment binding sites with engineered and wild-type heparanase show minimal overlap of fragments binding to mutation sites (Supplementary Figure 6). This supports the use of the engineered heparanase variant for high-throughput drug discovery efforts targeting heparanase.

In this study, we successfully applied a fragment-based drug discovery approach to identify novel chemical space for targeting heparanase, a critical enzyme linked to cancer and other diseases. Through a combination of crystallographic and computational techniques, we discovered a total of 33 fragments that bind to heparanase, with seven demonstrating inhibitory activity. The active fragments were dispersed amongst two active site sub-pockets and various allosteric pockets, which provides robust and diverse starting points to employ fragment modification strategies to develop a highly specific heparanase inhibitor. Notably, our fragment elaboration led to a seven-fold increase in potency for one of these fragments, highlighting the potential of this approach to uncover potent inhibitors. These findings not only expand the chemical diversity available for targeting heparanase but also provide a strong foundation for future drug development efforts aimed at combating heparanase-related diseases.

## Author Contributions

CJJ conceptualised research; LJD, HK, CW, and RLF performed experimental work; LJD, CJJ, and RLF performed data analysis; CN, CJJ, and RLF supervised research; LJD, CJJ, and RLF wrote the manuscript with input from all authors.

## Supporting Information

Additional data figures, tables of chemical structures, and methods, including crystallographic statistics.

## Acknowledgements

This research was undertaken in part using the MX1 and MX2 beamlines at the Australian Synchrotron, part of ANSTO, and made use of the Australian Cancer Research Foundation (ACRF) detector. Financial support by the Australian Research Council (ARC) Centre of Excellence for Innovations in Peptide and Protein Science (CE200100012) and the ARC Centre of Excellence for Innovations in Synthetic Biology (CE200100029) is gratefully acknowledged. CN thanks the ARC for a Discovery Project (DP230100079) and a Future Fellowship (FT220100010). LJD thanks the Medical Advances Without Animals trust for a PhD scholarship. The Biopolymer and Polymer Facility at the Research School of Chemistry is acknowledged for training and support.

