## Supplementary data and methods for "Fragment Screening and Structure-Guided Development of Heparanase Inhibitors Reveals Orthosteric and Allosteric Inhibition"

**Supplementary Table 1:** Crystallography statistics for the first 16 of the 32 apo heparanase structures used in Pan-density Dataset analysis

| **Structure ID** | Apo 1 | Apo 2 | Apo 3 | Apo 4 | Apo 5 | Apo 6 | Apo 7 | Apo 8 | Apo 9 | Apo 10 | Apo 11 | Apo 12 | Apo 13 | Apo 14 | Apo 15 | Apo 16 |
|---|---|---|---|---|---|---|---|---|---|---|---|---|---|---|---|---|
| **Data collection** | | | | | | | | | | | | | | | | |
| Space group | P $2_1$ $2_1$ $2_1$ | P $2_1$ $2_1$ $2_1$ | P $2_1$ $2_1$ $2_1$ | P $2_1$ $2_1$ $2_1$ | P $2_1$ $2_1$ $2_1$ | P $2_1$ $2_1$ $2_1$ | P $2_1$ $2_1$ $2_1$ | P $2_1$ $2_1$ $2_1$ | P $2_1$ $2_1$ $2_1$ | P $2_1$ $2_1$ $2_1$ | P $2_1$ $2_1$ $2_1$ | P $2_1$ $2_1$ $2_1$ | P $2_1$ $2_1$ $2_1$ | P $2_1$ $2_1$ $2_1$ | P $2_1$ $2_1$ $2_1$ | P $2_1$ $2_1$ $2_1$ |
| Cell Dimensions | | | | | | | | | | | | | | | | |
| *a, b, c (Å)* | 59.9 74.9 124.1 | 59.4 75.2 123.9 | 59.6 75.2 124.2 | 60.2 74.2 123.9 | 59.7 74.7 124.0 | 59.1 75.4 124.1 | 60.4 74.2 124.0 | 59.6 75.1 124.0 | 60.2 74.6 124.1 | 59.5 75.4 124.2 | 59.1 75.6 124.1 | 59.1 75.7 124.4 | 60.2 74.6 124.1 | 59.0 75.4 124.4 | 59.0 75.5 124.2 | 59.2 75.4 124.1 |
| α, β, γ (°) | 90 90 90 | 90 90 90 | 90 90 90 | 90 90 90 | 90 90 90 | 90 90 90 | 90 90 90 | 90 90 90 | 90 90 90 | 90 90 90 | 90 90 90 | 90 90 90 | 90 90 90 | 90 90 90 | 90 90 90 | 90 90 90 |
| Resolution (*Å*)* | 2.28 (2.28) | 1.68 (1.68) | 1.58 (1.58) | 1.78 (1.78) | 1.65 (1.65) | 1.61 (1.61) | 1.71 (1.71) | 1.66 (1.66) | 1.77 (1.77) | 1.70 (1.70) | 1.64 (1.64) | 1.73 (1.73) | 1.81 (1.81) | 1.69 (1.69) | 1.87 (1.87) | 1.63 (1.63) |
| $R_{merge}$ | 0.4499 (0.458) | 0.2014 (1.205) | 0.3263 (1.515) | 0.5323 (1.066 | 0.2829 (1.537) | 0.4302 (1.405) | 0.2264 (1.394) | 0.2243 (1.272) | 0.3108 (1.073) | 0.2595 (1.37) | 0.3388 (1.313) | 0.4004 (1.481) | 0.2089 (1.054) | 0.1747 (1.379) | 0.2442 (1.647) | 0.2134 (1.22) |
| $R_{pim}$ | 0.128 (0.129) | 0.05631 (0.330) | 0.09151 (0.429) | 0.1498 (0.299) | 0.07893 (0.423) | 0.1211 (0.398) | 0.06323 (0.379) | 0.06271 (0.344) | 0.08737 (0.288) | 0.07319 (0.375) | 0.09554 (0.364) | 0.1127 (0.405) | 0.05822 (0.284) | 0.04861 (0.377) | 0.06881 (0.457) | 0.05982 (0.341) |
| I/σI | 22.55 (5.09) | 14.39 (0.94) | 13.63 (0.88) | 10.90 (0.87) | 13.78 (0.87) | 12.87 (0.93) | 13.30 (0.89) | 13.47 (0.91) | 12.77 (0.82) | 12.37 (0.82) | 12.36 (0.90) | 10.35 (0.81) | 10.94 (0.87) | 11.83 (0.90) | 9.06 (0.70) | 13.60 (0.90) |
| $CC_{1/2}$ | 0.864 (0.886) | 0.993 (0.862) | 0.962 (0.742) | 0.842 (0.711) | 0.987 (0.735) | 0.949 (0.687) | 0.993 (0.823) | 0.991 (0.83) | 0.979 (0.718) | 0.985 (0.779) | 0.965 (0.742) | 0.95 (0.691) | 0.996 (0.797) | 0.997 (0.793) | 0.991 (0.708) | 0.992 (0.821) |
| Completeness (%) | 98.3 (83.6) | 99.9 (99.2) | 99.1 (91.2) | 99.3 (93.7) | 99.1 (91.6) | 99.3 (93.4) | 98.9 (91.2) | 99.1 (92.2) | 99.5 (94.8) | 99.2 (92.2) | 99.9 (99.9) | 99.5 (95.2) | 99.3 (94.1) | 99.9 (99.3) | 98.9 (88.6) | 99.9 (99.8) |
| Redundancy | 13.3 (12.6) | 13.7 (14.1) | 13.7 (13.3) | 13.6 (13.9) | 13.7 (14.0) | 13.7 (13.2) | 13.7 (14.1) | 13.8 (14.2) | 13.6 (14.0) | 13.7 (14.1) | 13.7 (13.8) | 13.7 (14.1) | 13.6 (14.0) | 13.7 (14.1) | 13.6 (13.8) | 13.7 (13.5) |
| **Refinement** | | | | | | | | | | | | | | | | |
| Resolution (*Å*) | 2.28 (2.28) | 1.68 (1.68) | 1.58 (1.58) | 1.78 (1.78) | 1.65 (1.65) | 1.61 (1.61) | 1.71 (1.71) | 1.66 (1.66) | 1.77 (1.77) | 1.70 (1.70) | 1.64 (1.64) | 1.73 (1.73) | 1.81 (1.81) | 1.69 (1.69) | 1.87 (1.87) | 1.63 (1.63) |
| No. reflections | 25697 (2128) | 64047 (6295) | 76182 (6913) | 53625 (4986) | 66807 (6080) | 72125 (6706) | 60373 (5508) | 65891 (6050) | 54950 (5132) | 61639 (5659) | 68935 (6788 | 58510 (5494) | 51001 (4760) | 62911 (6155) | 45901 (4058) | 69943 (6897) |
| $R_{work}$ | 0.1622 (0.216) | 0.1703 (0.279) | 0.1750 (0.283) | 0.1780 (0.268) | 0.1767 (0.304) | 0.1753 (0.280) | 0.1714 (0.283) | 0.1723 (0.277) | 0.1729 (0.330) | 0.1711 (0.275) | 0.1730 (0.261) | 0.1723 (0.270) | 0.1724 (0.316) | 0.1738 (0.286) | 0.1748 (0.292) | 0.1728 (0.275) |
| $R_{free}$ | 0.1945 (0.267) | 0.1897 (0.309) | 0.1904 (0.276) | 0.2061 (0.263) | 0.1939 (0.298) | 0.1927 (0.293) | 0.1947 (0.289) | 0.1898 (0.272) | 0.1943 (0.329) | 0.1894 (0.283) | 0.1915 (0.284) | 0.1918 (0.275) | 0.1961 (0.302) | 0.1964 (0.314) | 0.2009 (0.271) | 0.1896 (0.291) |
| No. atoms | 4112 | 4112 | 4112 | 4112 | 4112 | 4112 | 4112 | 4112 | 4112 | 4112 | 4112 | 4112 | 4112 | 4112 | 4112 | 4112 |
| Protein | 3761 | 3761 | 3761 | 3761 | 3761 | 3761 | 3761 | 3761 | 3761 | 3761 | 3761 | 3761 | 3761 | 3761 | 3761 | 3761 |
| Ligand/ion | 49 | 49 | 49 | 49 | 49 | 49 | 49 | 49 | 49 | 49 | 49 | 49 | 49 | 49 | 49 | 49 |
| Water | 302 | 302 | 302 | 302 | 302 | 302 | 302 | 302 | 302 | 302 | 302 | 302 | 302 | 302 | 302 | 302 |
| B-factors (overall) | 32.38 | 29.94 | 25.99 | 4112 | 29.07 | 25.6 | 31.1 | 27.65 | 31.22 | 30.1 | 26.24 | 27.27 | 34.46 | 29.5 | 33.66 | 28.48 |
| Protein | 31.29 | 28.7 | 24.73 | 3761 | 27.82 | 24.33 | 29.83 | 26.37 | 29.95 | 28.83 | 24.95 | 25.98 | 33.16 | 28.18 | 32.42 | 27.2 |
| Ligand/ion | 70.48 | 67.61 | 62.39 | 49 | 66.3 | 61.49 | 68.17 | 64.27 | 68.56 | 66.17 | 63.9 | 62.31 | 72.59 | 65.21 | 67.53 | 65.19 |
| Water | 39.81 | 39.35 | 35.78 | 302 | 38.7 | 35.56 | 40.89 | 37.68 | 41.06 | 40.01 | 36.17 | 37.67 | 44.57 | 40.15 | 43.69 | 38.5 |
| R.M.S. deviations | | | | | | | | | | | | | | | | |
| Bond lengths (*Å*) | 0.014 | 0.015 | 0.015 | 0.016 | 0.015 | 0.015 | 0.015 | 0.015 | 0.015 | 0.016 | 0.015 | 0.015 | 0.015 | 0.016 | 0.014 | 0.015 |
| Bond angles (°) | 1.91 | 1.92 | 1.87 | 1.95 | 1.87 | 1.86 | 1.92 | 1.92 | 1.91 | 1.92 | 1.86 | 1.9 | 1.9 | 1.88 | 1.87 | 1.86 |

* Statistics for the highest resolution shell are shown in parentheses

**Supplementary Table 1 (continued):** Crystallography statistics for the second 16 of the 32 apo heparanase structures used in Pan-density Dataset analysis

| **Structure ID** | Apo 17 | Apo 18 | Apo 19 | Apo 20 | Apo 21 | Apo 22 | Apo 23 | Apo 24 | Apo 25 | Apo 26 | Apo 27 | Apo 28 | Apo 29 | Apo 30 | Apo 31 | Apo 32 |
|---|---|---|---|---|---|---|---|---|---|---|---|---|---|---|---|---|
| **Data Collection** | | | | | | | | | | | | | | | | |
| Space group | $P\ 2_1\ 2_1\ 2_1$ | $P\ 2_1\ 2_1\ 2_1$ | $P\ 2_1\ 2_1\ 2_1$ | $P\ 2_1\ 2_1\ 2_1$ | $P\ 2_1\ 2_1\ 2_1$ | $P\ 2_1\ 2_1\ 2_1$ | $P\ 2_1\ 2_1\ 2_1$ | $P\ 2_1\ 2_1\ 2_1$ | $P\ 2_1\ 2_1\ 2_1$ | $P\ 2_1\ 2_1\ 2_1$ | $P\ 2_1\ 2_1\ 2_1$ | $P\ 2_1\ 2_1\ 2_1$ | $P\ 2_1\ 2_1\ 2_1$ | $P\ 2_1\ 2_1\ 2_1$ | $P\ 2_1\ 2_1\ 2_1$ | $P\ 2_1\ 2_1\ 2_1$ |
| Cell Dimensions | | | | | | | | | | | | | | | | |
| *a, b, c (Å)* | 59.6 75.5 124.5 | 59.0 75.7 124.4 | 58.9 75.7 124.3 | 60.6 73.8 124.1 | 58.9 75.8 124.5 | 59.3 75.4 124.2 | 59.2 75.4 124.3 | 59.1 75.4 124.2 | 59.2 75.7 124.2 | 59.1 75.4 124.1 | 59.0 75.7 124.2 | 58.9 75.4 124.4 | 59.2 75.5 124.2 | 59.1 75.8 124.2 | 58.8 75.6 124.0 | 59.6 75.5 124.5 |
| α, β, γ (°) | 90 90 90 | 90 90 90 | 90 90 90 | 90 90 90 | 90 90 90 | 90 90 90 | 90 90 90 | 90 90 90 | 90 90 90 | 90 90 90 | 90 90 90 | 90 90 90 | 90 90 90 | 90 90 90 | 90 90 90 | 90 90 90 |
| Resolution (*Å*)* | 1.88 (1.88) | 2.21 (2.21) | 1.86 (1.86) | 1.66 (1.66) | 2.09 (2.09) | 1.86 (1.86) | 1.85 (1.85) | 2.03 (2.03) | 2.19 (2.19) | 1.66 (1.66) | 1.70 (1.70) | 1.63 (1.63) | 1.81 (1.81) | 2.36 (2.36) | 1.64 (1.64) | 1.72 (1.72) |
| $R_{merge}$ | 0.4443 (1.54) | 0.04455 (0.434) | 0.4161 (1.624) | 0.1762 (1.298) | 0.05719 (0.514) | 0.3689 (1.347) | 0.2203 (1.441) | 0.5299 (1.349) | 0.543 (1.369) | 0.4011 (1.391) | 0.03976 (0.328) | 0.3915 (1.277) | 0.537 (1.053) | 0.08399 (0.433) | 0.3029 (1.297) | 0.3574 (1.524) |
| $R_{pim}$ | 0.1252 (0.426) | 0.04455 (0.434) | 0.1174 (0.448) | 0.04951 (0.354) | 0.05719 (0.514) | 0.1059 (0.378) | 0.06204 (0.397) | 0.1498 (0.386) | 0.1533 (0.388) | 0.1132 (0.382) | 0.03976 (0.328) | 0.1101 (0.357) | 0.152 (0.303) | 0.08399 (0.433) | 0.08506 (0.359) | 0.1029 (0.423) |
| I/σI | 9.28 (0.87) | 10.47 (1.47) | 9.25 (0.78) | 15.80 (0.89) | 6.88 (1.20) | 11.98 (0.88) | 8.87 (0.75) | 7.69 (0.77) | 6.84 (0.81) | 10.13 (0.81) | 9.08 (1.44) | 11.14 (0.87) | 8.87 (0.81) | 7.03 (1.55) | 10.56 (0.84) | 10.06 (0.78) |
| $CC_{1/2}$ | 0.903 (0.611) | 0.998 (0.620) | 0.925 (0.635) | 0.995 (0.738) | 0.997 (0.560) | 0.955 (0.702) | 0.994 (0.736) | 0.917 (0.574) | 0.919 (0.605) | 0.947 (0.735) | 0.998 (0.87) | 0.945 (0.792) | 0.931 (0.768) | 0.989 (0.585) | 0.967 (0.765) | 0.950 (0.730) |
| Completeness (%) | 99.1 (90.9) | 98.8 (88.1) | 99.3 (93.0) | 99.7 (97.6) | 99.3 (93.4) | 99.4 (94.6) | 99.3 (93.8) | 99.9 (99.9) | 98.4 (90.5) | 99.5 (94.8) | 96.3 (66.7) | 99.6 (96.6) | 99.8 (98.2) | 99.6 (97.9) | 99.4 (94.4) | 98.4 (84.6) |
| Redundancy | 13.6 (13.8) | 1.9 (1.9) | 13.6 (13.8) | 13.7 (14.1) | 1.9 (1.9) | 13.5 (13.8) | 13.6 (13.9) | 13.6 (13.1) | 13.8 (13.7) | 13.7 (14.1) | 1.9 (1.9) | 13.7 (13.5) | 13.6 (13.9) | 1.9 (1.9) | 13.7 (13.8) | 13.7 (14.0) |
| **Refinement** | | | | | | | | | | | | | | | | |
| Resolution (*Å*) | 1.88 (1.88) | 2.21 (2.21) | 1.86 (1.86) | 1.66 (1.66) | 2.09 (2.09) | 1.86 (1.86) | 1.85 (1.85) | 2.03 (2.03) | 2.19 (2.19) | 1.66 (1.66) | 1.70 (1.70) | 1.63 (1.63) | 1.81 (1.81) | 2.36 (2.36) | 1.64 (1.64) | 1.72 (1.72) |
| No. reflections | 46445 (4224) | 28067 (2467) | 47023 (4329) | 66349 (6392) | 33209 (3083) | 47365 (4437) | 47699 (4427) | 36725 (3630) | 28733 (2611) | 66975 (6261) | 59476 (4059) | 69813 (6696) | 51168 (4950) | 23487 (2260) | 68442 (6386) | 58534 (4974) |
| $R_{work}$ | 0.1734 (0.265) | 0.1739 (0.259) | 0.1728 (0.278) | 0.1716 (0.274) | 0.1814 (0.270) | 0.1718 (0.269) | 0.1728 (0.288) | 0.1729 (0.253) | 0.1774 (0.276) | 0.1734 (0.270) | 0.1867 (0.292) | 0.1776 (0.279) | 0.1865 (0.298) | 0.1743 (0.247) | 0.1730 (0.270) | 0.1735 (0.285) |
| $R_{free}$ | 0.1986 (0.260) | 0.2076 (0.289) | 0.1969 (0.260) | 0.1884 (0.278) | 0.2160 (0.275) | 0.1945 (0.268) | 0.1981 (0.289) | 0.2033 (0.279) | 0.2117 (0.305) | 0.1924 (0.290) | 0.2075 (0.306) | 0.1964 (0.294) | 0.2066 (0.304) | 0.2080 (0.278) | 0.1927 (0.279) | 0.1964 (0.295) |
| No. atoms | 4112 | 4112 | 4112 | 4112 | 4112 | 4112 | 4112 | 4112 | 4112 | 4112 | 4112 | 4112 | 4112 | 4112 | 4112 | 4112 |
| Protein | 3761 | 3761 | 3761 | 3761 | 3761 | 3761 | 3761 | 3761 | 3761 | 3761 | 3761 | 3761 | 3761 | 3761 | 3761 | 3761 |
| Ligand/ion | 49 | 49 | 49 | 49 | 49 | 49 | 49 | 49 | 49 | 49 | 49 | 49 | 49 | 49 | 49 | 49 |
| Water | 302 | 302 | 302 | 302 | 302 | 302 | 302 | 302 | 302 | 302 | 302 | 302 | 302 | 302 | 302 | 302 |
| B-factors (overall) | 26.8 | 36.98 | 28.98 | 30.74 | 31.18 | 33.18 | 30.94 | 32.27 | 36 | 24.79 | 21.47 | 25.2 | 27.27 | 27.01 | 25.35 | 28.13 |
| Protein | 25.54 | 35.98 | 27.73 | 29.47 | 29.95 | 31.89 | 29.62 | 30.84 | 34.77 | 23.54 | 20.33 | 24.03 | 26.03 | 26.04 | 24.09 | 26.91 |
| Ligand/ion | 63.24 | 73.14 | 64.8 | 66.33 | 68.9 | 69.85 | 66.37 | 69.09 | 69.87 | 61.96 | 57.06 | 60.5 | 63.65 | 62.13 | 62.31 | 64.13 |
| Water | 36.54 | 43.52 | 38.63 | 40.77 | 40.38 | 43.21 | 41.57 | 44.16 | 45.74 | 34.32 | 29.82 | 34.08 | 36.85 | 33.43 | 35.08 | 37.5 |
| R.M.S. deviations | | | | | | | | | | | | | | | | |
| Bond lengths (*Å*) | 0.015 | 0.014 | 0.015 | 0.015 | 0.014 | 0.015 | 0.015 | 0.014 | 0.014 | 0.015 | 0.015 | 0.015 | 0.015 | 0.014 | 0.015 | 0.015 |
| Bond angles (°) | 1.92 | 1.91 | 1.89 | 1.87 | 1.88 | 1.9 | 1.87 | 1.88 | 1.88 | 1.9 | 1.87 | 1.92 | 1.92 | 1.85 | 1.91 | 1.9 |

* Statistics for the highest resolution shell are shown in parentheses

**Supplementary Table 2:** Crystallography statistics for heparanase bound to fragments from the JBS:Frag Xtal Screen

| **Structure ID** | J4 (**3**) | J11 (**4**) | J13 (**20**) | J16 (**15**) | J21 (**18**) | J25 (**5**) | J28 (**21**) | J29 (**6**) | J31 (**19**) | J32 (**7**) | J34 (**8**) | J36 (**9**) | J37 (**10**) |
|---|---|---|---|---|---|---|---|---|---|---|---|---|---|
| **PDB ID** | 9O1R | 9O1S | 9O1T | 9O1U | 9O1V | 9O1W | 9O1X | 9O1Y | 9O1Z | 9O20 | 9O21 | 9O22 | 9O23 |
| **Data collection** | | | | | | | | | | | | | |
| Space group | $P 2_1 2_1 2_1$ | $P 2_1 2_1 2_1$ | $P 2_1 2_1 2_1$ | $P 2_1 2_1 2_1$ | $P 2_1 2_1 2_1$ | $P 2_1 2_1 2_1$ | $P 2_1 2_1 2_1$ | $P 2_1 2_1 2_1$ | $P 2_1 2_1 2_1$ | $P 2_1 2_1 2_1$ | $P 2_1 2_1 2_1$ | $P 2_1 2_1 2_1$ | $P 2_1 2_1 2_1$ |
| Cell dimensions | | | | | | | | | | | | | |
| *a, b, c (Å)* | 59.1 75.8 125.4 | 58.6 75.0 125.1 | 59.2 76.0 126.3 | 59.4 66.5 122.5 | 58.7 74.7 124.1 | 58.4 75.2 124.2 | 58.7 74.5 123.8 | 58.7 74.7 125.1 | 58.5 74.6 15.1 | 58.9 74.5 124.1 | 58.8 74.9 124.2 | 59.4 76.2 125.2 | 58.8 74.9 124.9 |
| a, b, c (°) | 90 90 90 | 90 90 90 | 90 90 90 | 90 90 90 | 90 90 90 | 90 90 90 | 90 90 90 | 90 90 90 | 90 90 90 | 90 90 90 | 90 90 90 | 90 90 90 | 90 90 90 |
| Resolution (*Å*)* | 48.31-1.89 (1.94-1.89) | 46.20-1.80 (1.84-1.80) | 48.56-1.90 (1.94-1.9) | 45.03-1.90 (1.97-1.9) | 46.15-2.05 (2.11-2.05) | 46.10-1.90 (1.94-1.90) | 47.63-1.90 (1.94-1.90) | 47.93-2.00 (2.05-2.00) | 47.94-2.50 (2.60-2.50) | 47.70-1.83 (1.87-1.83) | 47.81-1.82 (1.86-1.82) | 46.81-2.10 (2.16-2.10) | 47.98-2.20 (2.28 - 2.20) |
| $R_{merge}$ | 0.253 (4.874) | 0.893 (44.091) | 0.154 (6.870) | 0.482 (3.732) | 0.538 (17.174) | 0.818 (15.874) | 0.380 (3.205) | 1.551 (61.493) | 0.805 (9.727) | 0.607 (15.722) | 0.319 (6.041) | 0.582 (10.716) | 0.245 (1.444) |
| $R_{pim}$ | 0.101 (2.018) | 0.249 (12.213) | 0.043 (2.003) | 0.135 (1.030) | 0.154 (4.876) | 0.238 (5.030) | 0.106 (0.875) | 0.451 (19.235) | 0.214 (2.709) | 0.170 (4.336) | 0.090 (1.670) | 0.163 (3.068) | 0.133 (0.794) |
| $I/\sigma I$ | 10.80 (2.20) | 6.60 (0.50) | 12.8 (0.80) | 5.40 (0.90) | 6.50 (0.70) | 12.70 (2.60) | 6.90 (0.90) | 5.10 (0.60) | 4.90 (1.00) | 5.90 (0.50) | 6.80 (0.70) | 9.90 (1.90) | 3.40 (0.90) |
| $CC_{1/2}$ | 0.996 (0.316) | 0.991 (0.423) | 0.999 (0.570) | 0.986 (0.402) | 0.984 (0.363) | 0.829 (0.470) | 0.991 (0.400) | 0.861 (0.643) | 0.983 (0.344) | 0.994 (0.303) | 0.995 (0.302) | 0.992 (0.610) | 0.980 (0.344) |
| Completeness (%) | 98.7 (91.3) | 99.8 (98.9) | 100.0 (100.0) | 100.0 (100.0) | 99.9 (100.0) | 99.9 (99.4) | 99.8 (99.8) | 99.9 (99.4) | 99.7 (99.3) | 100.0 (100.0) | 100.0 (100.0) | 100.0 (100.0) | 97.7 (98.5) |
| Redundancy | 13.7 (13.0) | 13.6 (13.9) | 13.6 (12.7) | 13.5 (13.9) | 13.5 (13.3) | 13.4 (12.6) | 13.7 (14.1) | 12.8 (11.4) | 13.3 (13.6) | 13.5 (13.8) | 13.5 (13.8) | 13.6 (13.0) | 4.2 (4.2) |
| **Refinement** | | | | | | | | | | | | | |
| Resolution (*Å*) | 48.31-1.89 (1.94-1.89) | 46.20-1.80 (1.84-1.80) | 48.56-1.90 (1.94-1.9) | 45.03-1.90 (1.97-1.9) | 46.15-2.05 (2.11-2.05) | 46.10-1.90 (1.94-1.90) | 47.63-1.90 (1.94-1.90) | 47.93-2.00 (2.05-2.00) | 47.94-2.50 (2.60-2.50) | 47.70-1.83 (1.87-1.83) | 47.81-1.82 (1.86-1.82) | 46.81-2.10 (2.16-2.10) | 47.98-2.20 (2.28 - 2.20) |
| No. reflections | 44674 (22562) | 51239 (2492) | 45669 (2793) | 38916 (2742) | 34589 (2639) | 43220 (2691) | 43410 (2822) | 37301 (2293) | 19527 (2727) | 48923 (2663) | 49906 (2715) | 33825 (2745) | 28670 (2586) |
| $R_{work}$ | 0.1733 | 0.1866 | 0.1848 | 0.1094 | 0.2204 | 0.1706 | 0.1861 | 0.2019 | 0.1960 | 0.1969 | 0.1894 | 0.1766 | 0.1866 |
| $R_{free}$ | 0.2139 | 0.2255 | 0.2158 | 0.2475 | 0.2702 | 0.2075 | 0.2341 | 0.2363 | 0.2440 | 0.2333 | 0.2228 | 0.2250 | 0.2377 |
| No. atoms | 3992 | 4030 | 4053 | 4108 | 3753 | 4133 | 4230 | 3849 | 3786 | 3960 | 3999 | 3948 | 3922 |
| Protein | 3625 | 3623 | 3645 | 3651 | 3620 | 3641 | 3668 | 3606 | 3629 | 3629 | 3628 | 3663 | 3631 |
| Ligand/ion | 39 | 66 | 60 | 157 | 48 | 80 | 104 | 37 | 43 | 40 | 58 | 37 | 48 |
| Water | 328 | 341 | 348 | 300 | 85 | 412 | 458 | 206 | 114 | 291 | 313 | 248 | 243 |
| B-factors (overall) | 31.29 | 31.33 | 36.60 | 20.55 | 46.06 | 27.07 | 24.90 | 32.78 | 44.25 | 29.36 | 29.57 | 459 | 458 |
| Protein | 30.35 | 30.09 | 35.66 | 19.53 | 45.98 | 25.72 | 23.24 | 32.34 | 43.87 | 28.58 | 28.50 | 35.52 | 34.49 |
| Ligand/ion | 55.08 | 56.70 | 58.49 | 30.11 | 53.87 | 43.25 | 47.00 | 47.40 | 82.40 | 49.85 | 47.76 | 34.88 | 33.82 |
| Water | 38.83 | 39.57 | 42.67 | 28.00 | 45.08 | 35.91 | 33.19 | 37.89 | 41.93 | 36.31 | 38.56 | 56.33 | 61.73 |
| R.M.S. deviations | | | | | | | | | | | | | |
| Bond lengths (*Å*) | 0.006 | 0.007 | 0.008 | 0.006 | 0.003 | 0.006 | 0.006 | 0.007 | 0.002 | 0.006 | 0.006 | 0.007 | 0.007 |
| Bond angles (*Å*) | 0.842 | 0.856 | 0.896 | 0.906 | 0.588 | 0.849 | 0.855 | 0.911 | 0.506 | 0.851 | 0.855 | 1.245 | 0.858 |

* Statistics for the highest resolution shell are shown in parentheses

**Supplementary Table 2 (cont.):** Crystallography statistics for heparanase bound to fragments from the JBS:Frag Xtal Screen

| **Structure ID** | J40 (**11**) | J41 (**12**) | J51 (**13**) | J56 (**22**) | J59 (**23**) | J61 (**24**) | J69 (**25**) | J71 (**14**) | J72 (**16**) | J74 (**17**) | J82 (**26**) | J85 (**1**) | J91 (**27**) | J92 (**2**) |
|---|---|---|---|---|---|---|---|---|---|---|---|---|---|---|
| **PDB ID** | 9O24 | 9O25 | 9O26 | 9O27 | 9O28 | 9O29 | 9O2A | 9O2B | 9O2C | 9O2D | 9O2E | 9O2F | 9O2G | 9O2H |
| **Data collection** | | | | | | | | | | | | | | |
| Space group | $P 2_1 2_1 2_1$ | $P 2_1 2_1 2_1$ | $P 2_1 2_1 2_1$ | $P 2_1 2_1 2_1$ | $P 2_1 2_1 2_1$ | $P 2_1 2_1 2_1$ | $P 2_1 2_1 2_1$ | $P 2_1 2_1 2_1$ | $P 2_1 2_1 2_1$ | $P 2_1 2_1 2_1$ | $P 2_1 2_1 2_1$ | $P 2_1 2_1 2_1$ | $P 2_1 2_1 2_1$ | $P 2_1 2_1 2_1$ |
| Cell dimensions | | | | | | | | | | | | | | |
| *a, b, c (Å)* | 58.7 74.6 124.3 | 58.8 74.7 124.4 | 57.9 73.8 124.1 | 58.4 73.9 123.5 | 58.8 74.9 124.8 | 59.5 74.0 124.9 | 58.3 74.8 124.3 | 59.0 75.0 125.3 | 59.3 75.6 125.4 | 58.6 74.2 123.4 | 58.9 75.1 124.9 | 59.2 74.7 124.9 | 58.2 73.9 123.6 | 58.4 74.0 123.9 |
| a, b, c (°) | 90 90 90 | 90 90 90 | 90 90 90 | 90 90 90 | 90 90 90 | 90 90 90 | 90 90 90 | 90 90 90 | 90 90 90 | 90 90 90 | 90 90 90 | 90 90 90 | 90 90 90 | 90 90 90 |
| Resolution (*Å*)* | 46.13-2.60 (2.72-2.60) | 47.80-1.65 (1.68-1.65) | 45.60-2.35 (2.43-2.35) | 47.40-2.10 (2.16-2.10) | 46.30-1.81 (1.85-1.81) | 46.40-2.20 (2.27-2.20) | 42.54-2.10 (2.16-2.10) | 46.39-2.20 (2.27-2.20) | 46.66-1.70 (1.73-1.70) | 47.44-2.30 (2.38-2.30) | 48.02-2.10 (2.16-2.10) | 46.41-2.50 (2.60-2.50) | 45.73-2.28 (2.36-2.28) | 47.52-1.75 (1.78-1.75) |
| $R_{merge}$ | 0.922 (10.339) | 0.333 (16.111) | 1.527 (30.053) | 0.433 (6.759) | 0.370 (9.673) | 0.432 (4.927) | 0.147 (1.136) | 0.458 (5.684) | 0.174 (8.546) | 1.011 (10.351) | 0.347 (3.147) | 0.292 (2.113) | 0.568 (9.419) | 0.252 (11.110) |
| $R_{pim}$ | 0.287 (3.063) | 0.094 (4.499) | 0.436 (8.797) | 0.122 (1.913) | 0.103 (2.651) | 0.120 (1.342) | 0.071 (0.551) | 0.132 (1.665) | 0.049 (2.367) | 0.289 (3.072) | 0.101 (0.945) | 0.132 (0.959) | 0.158 (2.559) | 0.071 (3.052) |
| $I/\sigma I$ | 7.70 (2.90) | 7.60 (0.70) | 5.40 (0.90) | 7.70 (1.30) | 7.80 (1.10) | 6.00 (0.90) | 8.20 (1.50) | 6.00 (1.40) | 8.40 (0.50) | 4.40 (0.80) | 8.60 (1.90) | 5.40 (0.90) | 6.60 (0.90) | 8.80 (0.90) |
| $CC_{1/2}$ | 0.877 (0.460) | 0.997 (0.383) | 0.944 (0.482) | 0.992 (0.302) | 0.996 (0.545) | 0.991 (0.312) | 0.987 (0.596) | 0.988 (0.406) | 0.999 (0.373) | 0.960 (0.291) | 0.992 (0.684) | 0.986 (0.417) | 0.983 (0.379) | 0.998 (0.265) |
| Completeness (%) | 100.0 (100.0) | 100.0 (100.0) | 100.0 (100.0) | 100.0 (100.0) | 99.3 (97.9) | 100.0 (100.0) | 82.7 (83.7) | 99.7 (99.5) | 100.0 (100.0) | 100.0 (100.0) | 98.0 (88.4) | 99.8 (99.8) | 99.0 (98.0) | 100.0 (100.0) |
| Redundancy | 13.2 (13.2) | 13.5 (13.7) | 13.1 (12.6) | 13.6 (13.5) | 13.7 (14.2) | 13.6 (14.2) | 5.2 (5.0) | 13.0 (12.1) | 13.6 (13.8) | 13.2 (12.1) | 12.8 (11.6) | 5.9 (5.8) | 13.7 (14.4) | 13.6 (13.9) |
| **Refinement** | | | | | | | | | | | | | | |
| Resolution (*Å*) | 46.13-2.60 (2.72-2.60) | 47.80- 1.65 (1.68-1.65) | 45.60-2.35 (2.43-2.35) | 47.40-2.10 (2.16-2.10) | 46.30-1.81 (1.85-1.81) | 46.40-2.20 (2.27-2.20) | 42.54-2.10 (2.16-2.10) | 46.39-2.20 (2.27-2.20) | 46.66-1.70 (1.73-1.70) | 47.44-2.30 (2.38-2.30) | 48.02-2.10 (2.16-2.10) | 46.41-2.50 (2.60-2.50) | 45.73-2.28 (2.36-2.28) | 47.52-1.75 (1.78-1.75) |
| No. reflections | 17303 (2732) | 66851 (2732) | 22705 (2696) | 31756 (2842) | 50644 (2580) | 28743 (2841) | 26622 (3005) | 28860 (2829) | 62656 (2773) | 24588 (3011) | 31561 (2542) | 19781 (2740) | 24653 (2675) | 54896 (2832) |
| $R_{work}$ | 0.1673 | 0.1830 | 0.1983 | 0.1833 | 0.1778 | 0.2026 | 0.2133 | 0.1877 | 0.1937 | 0.1970 | 0.2349 | 0.2088 | 0.1944 | 0.1836 |
| $R_{free}$ | 0.2361 | 0.2165 | 0.2340 | 0.2225 | 0.2095 | 0.2404 | 0.2607 | 0.2327 | 0.2152 | 0.2522 | 0.2822 | 0.2661 | 0.2447 | 0.2107 |
| No. atoms | 3748 | 3972 | 3757 | 3852 | 4038 | 3818 | 3764 | 3834 | 3881 | 3927 | 3852 | 3749 | 3730 | 3952 |
| Protein | 3620 | 3636 | 3620 | 3619 | 3630 | 3620 | 3620 | 3628 | 3614 | 3626 | 3611 | 3620 | 3609 | 3608 |
| Ligand/ion | 29 | 36 | 30 | 26 | 58 | 39 | 26 | 57 | 54 | 51 | 21 | 54 | 32 | 55 |
| Water | 99 | 300 | 107 | 207 | 350 | 159 | 118 | 149 | 213 | 250 | 22 | 75 | 89 | 289 |
| B-factors (overall) | 31.57 | 29.29 | 41.26 | 36.42 | 28.90 | 39.29 | 37.25 | 40.10 | 37.16 | 30.62 | 23.95 | 42.95 | 44.64 | 32.61 |
| Protein | 31.43 | 28.48 | 41.10 | 36.00 | 27.83 | 39.00 | 37.10 | 39.65 | 36.69 | 30.29 | 23.69 | 42.72 | 44.48 | 31.89 |
| Ligand/ion | 48.69 | 49.06 | 61.60 | 52.65 | 48.21 | 57.53 | 46.43 | 59.98 | 52.98 | 41.09 | 29.72 | 61.10 | 71.74 | 45.75 |
| Water | 31.67 | 36.72 | 40.91 | 41.67 | 36.88 | 41.46 | 39.80 | 43.59 | 41.12 | 33.26 | 27.63 | 41.32 | 44.68 | 39.46 |
| R.M.S. deviations | | | | | | | | | | | | | | |
| Bond lengths (*Å*) | 0.008 | 0.006 | 0.002 | 0.007 | 0.006 | 0.007 | 0.002 | 0.003 | 0.006 | 0.004 | 0.007 | 0.003 | 0.008 | 0.006 |
| Bond angles (°) | 0.908 | 0.852 | 0.511 | 0.923 | 0.840 | 0.874 | 0.551 | 0.602 | 0.810 | 0.633 | 0.856 | 0.572 | 0.920 | 0.891 |

* Statistics for the highest resolution shell are shown in parentheses

**Supplementary Table 3:** Fragments identified to bind heparanase in crystallographic screening. *Denotes fragments bound in more than one binding site on heparanase.

| | | | | | |
|---|---|---|---|---|---|
| Active site (sub pocket 1) | 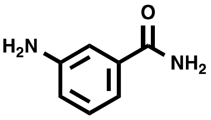 | **1** | | 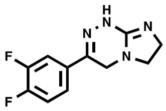 | **4*** |
| | 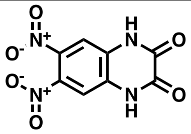 | **2** | Remote sites | 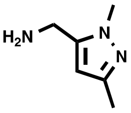 | **20** |
| Active site (sub pocket 2) | 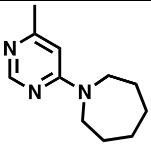 | **3** | | 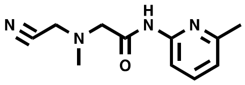 | **15*** |
| | 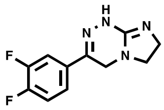 | **4*** | | 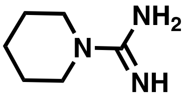 | **18** |
| | 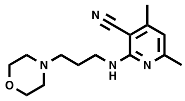 | **5** | | 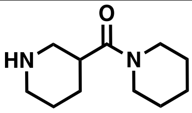 | **19** |
| | 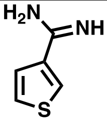 | **6** | | 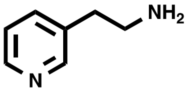 | **21** |
| | 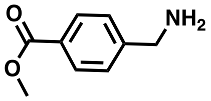 | **7** | | 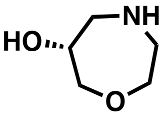 | **22** |
| | 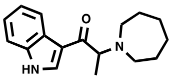 | **8** | | 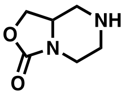 | **23** |
| | 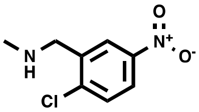 | **9** | | 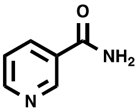 | **24*** |
| | 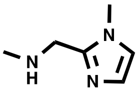 | **10** | | 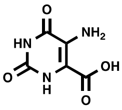 | **25** |
| | 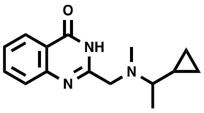 | **11** | | 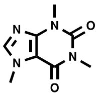 | **14*** |
| | 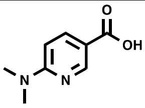 | **12** | | 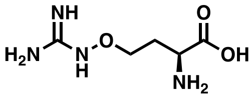 | **16*** |
| | 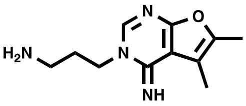 | **13** | | 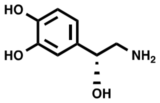 | **17** |
| | 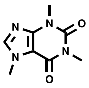 | **14*** | | 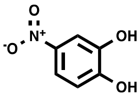 | **26** |
| | 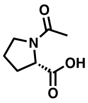 | **27** | | | |

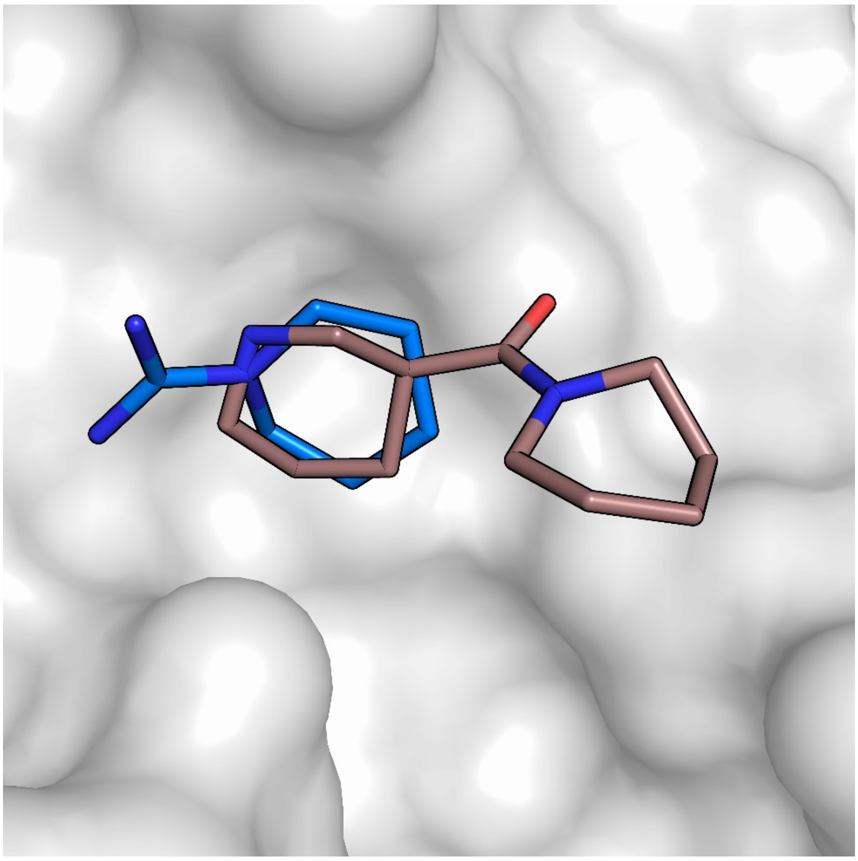

**Supplementary Figure 1:** The near-perfect overlap in the binding pose of the common moiety of **18** (blue) and **19** (brown) presents an opportunity for fragment merging.

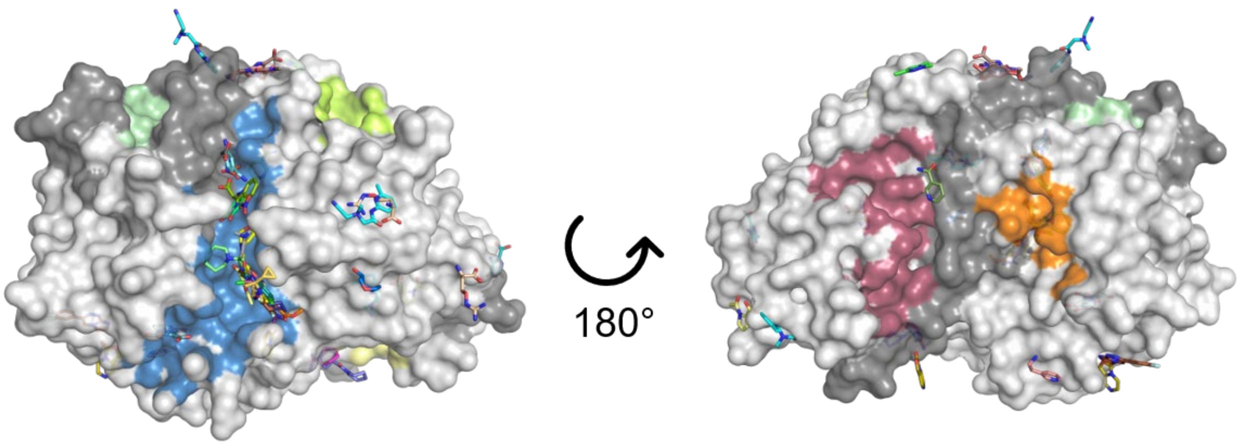

**Supplementary Figure 2:** SiteMap-predicted binding sites and their poor overlap with binding sites of fragments (coloured sticks) identified in the crystallographic fragment screen. Heparanase is shown as grey surface, with computationally-predicted binding sites as coloured surfaces. The active site is shown in blue.

**Supplementary Table 4:** Fragments selected from docking studies and their G-score for each site.

| Site | ID | G-Score (kcal/mol) | Chemical Structure | Site | ID | G-Score (kcal/mol) | Chemical Structure |
|---|---|---|---|---|---|---|---|
| Active site | **34** | -8.910 | 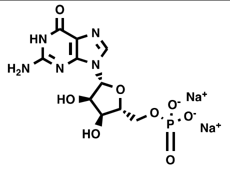 | Site B | **42** | -5.080 | 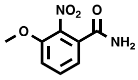 |
| | **29** | -8.560 | 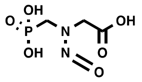 | | **28** | -4.870 | 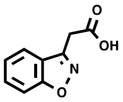 |
| | **35** | -8.080 | 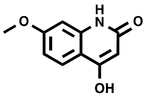 | | **43** | -4.830 | 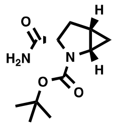 |
| | **36** | -7.810 | 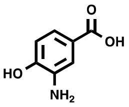 | | **44** | -4.780 | 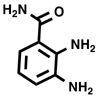 |
| | **31** | -7.735 | 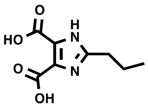 | Site C | **45** | -5.340 | 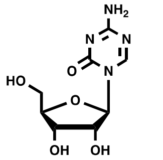 |
| Site A | **29** | -7.130 | 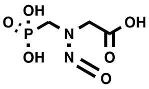 | | **39** | -5.184 | 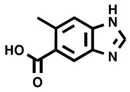 |
| | **37** | -6.670 | 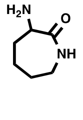 | Site D | **29** | -4.320 | 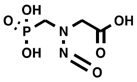 |
| | **38** | -6.460 | 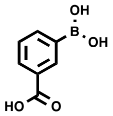 | Site E | **41** | -5.300 | 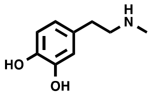 |
| | **39** | -6.440 | 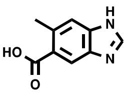 | | **46** | -5.290 | 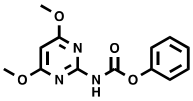 |
| | **40** | -6.330 | 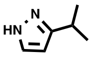 | | **30** | -4.450 | 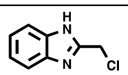 |
| Site B | **41** | -5.494 | 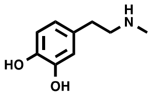 | | **47** | -4.358 | 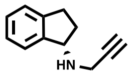 |

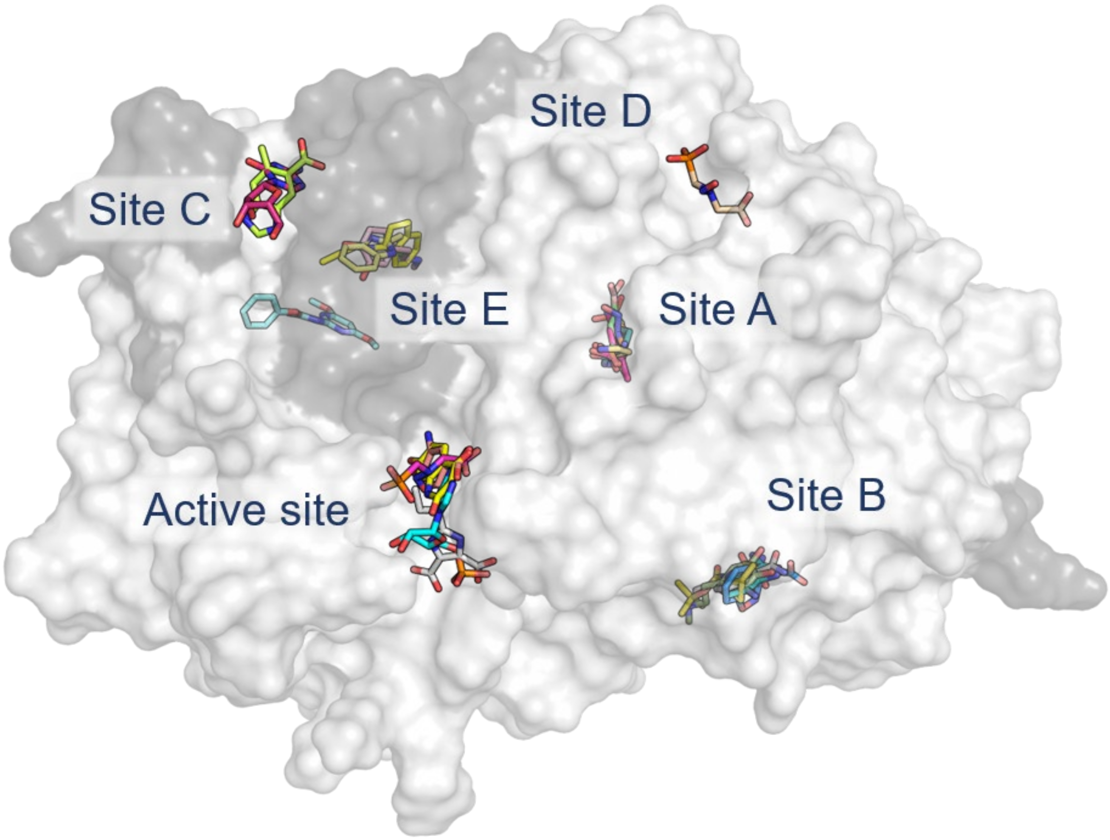

**Supplementary Figure 3: Docked poses of 18 fragments selected for experimental testing.** Heparanase is shown in grey surface, and fragments are shown in sticks.

**Supplementary Table 5:** Crystallography statistics for heparanase bound to fragments from computational studies.

| **Structure ID** | C2 (**29**) | C5 (**31**) | C12 (**28**) | C17 (**30**) |
|---|---|---|---|---|
| **PDB ID** | 9O2I | 9O2J | 9O2K | 9O2L |
| **Data collection** | | | | |
| Space group | P $2_1$ $2_1$ $2_1$ | P $2_1$ $2_1$ $2_1$ | P $2_1$ $2_1$ $2_1$ | P $2_1$ $2_1$ $2_1$ |
| Cell dimensions | | | | |
| *a, b, c (Å)* | 58.9 74.7 124.6 | 58.5 75.2 123.2 | 58.9 75.2 125.1 | 58.6 74.9 124.8 |
| a, b, c (°) | 90 90 90 | 90 90 90 | 90 90 90 | 90 90 90 |
| Resolution (*Å*)* | 47.83-2.50 (2.60-2.50) | 46.15-2.15 (2.22-2.15) | 48.07-1.73 (1.77 - 1.73) | 47.95 - 1.90 (1.94 - 1.90) |
| $R_{merge}$ | 0.603 (1.856) | 0.829 (47.325) | 0.627 (38.100) | 0.422 (7.350) |
| $R_{pim}$ | 0.174 (0.531) | 0.230 (12.940) | 0.177 (11.867) | 0.120 (2.147) |
| $I/\sigma I$ | 5.20 (1.60) | 6.30 (0.30) | 7.80 (0.70) | 6.30 (0.80) |
| $CC_{1/2}$ | 0.948 (0.351) | 0.990 (0.548) | 0.964 (0.428) | 0.993 (0.675) |
| Completeness (%) | 99.8 (98.9) | 100.0 (100.0) | 99.2 (86.0) | 100.0 (100.0) |
| Redundancy | 12.8 (13.0) | 13.6 (14.2) | 13.5 (11.4) | 13.4 (13.2) |
| **Refinement** | | | | |
| Resolution (*Å*) | 47.83-2.50 (2.60-2.50) | 46.15-2.15 (2.22-2.15) | 48.07-1.73 (1.77 - 1.73) | 47.95 -1.90 (1.94 - 1.90) |
| No. reflections | 19602 (2741) | 29464 (2559) | 57890 (2286) | 43752 (2562) |
| $R_{work}$ | 0.2179 | 0.2143 | 0.2059 | 0.1945 |
| $R_{free}$ | 0.2708 | 0.2600 | 0.2366 | 0.2316 |
| No. atoms | 3765 | 3720 | 3993 | 3936 |
| Protein | 3620 | 3637 | 3608 | 3661 |
| Ligand/ion | 17 | 19 | 82 | 42 |
| Water | 128 | 64 | 303 | 233 |
| B-factors (overall) | 21.60 | 47.72 | 27.69 | 32.33 |
| Protein | 31.62 | 47.62 | 36.75 | 31.69 |
| Ligand/ion | 34.18 | 63.99 | 43.7 | 58.50 |
| Water | 19.45 | 47.21 | 34.56 | 37.67 |
| R.m.s. deviations | | | | |
| Bond lengths (*Å)* | 0.002 | 0.008 | 0.006 | 0.006 |
| Bond angles (*Å*) | 0.507 | 0.964 | 0.846 | 0.816 |

* Statistics for the highest resolution shell are shown in parentheses

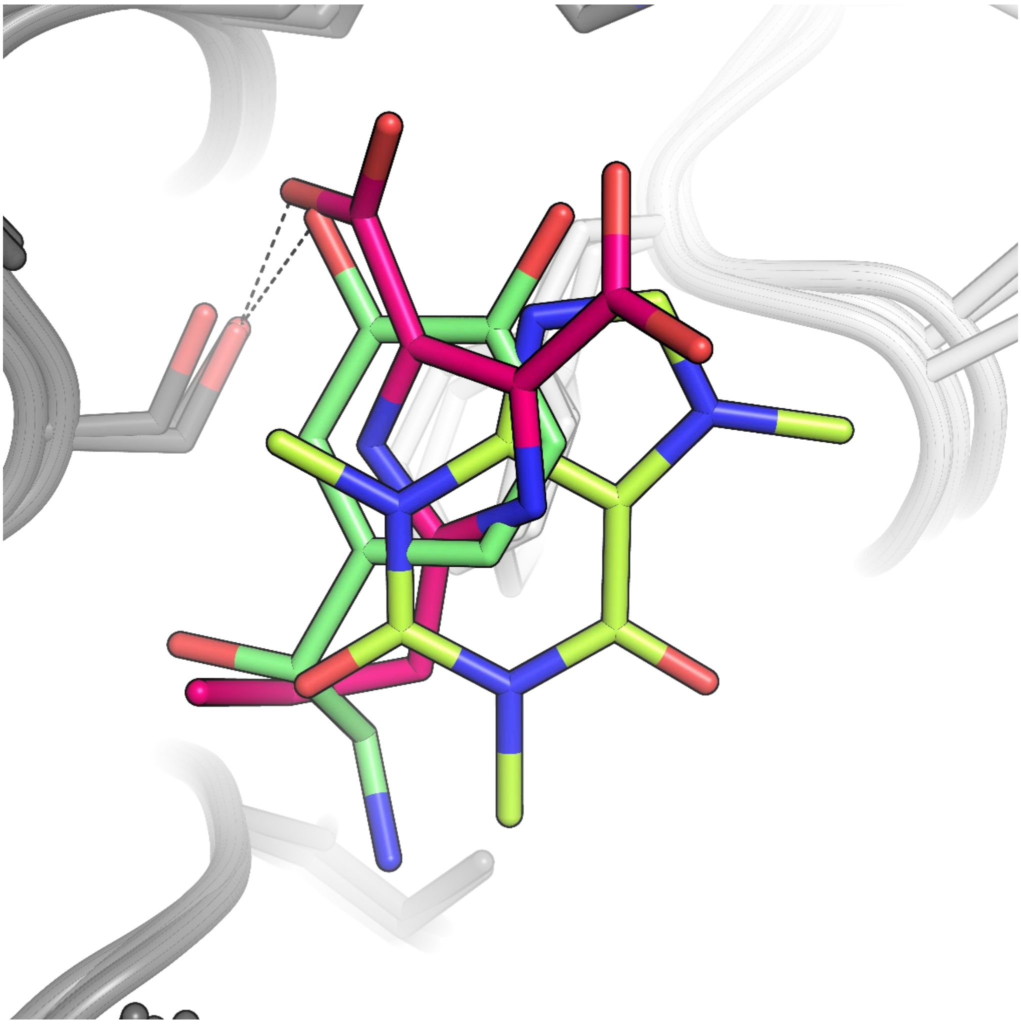

**Supplementary Figure 4:** The binding modes of **14** (yellow), **17** (green), and **31** (pink) colocalised at the same allosteric binding site.

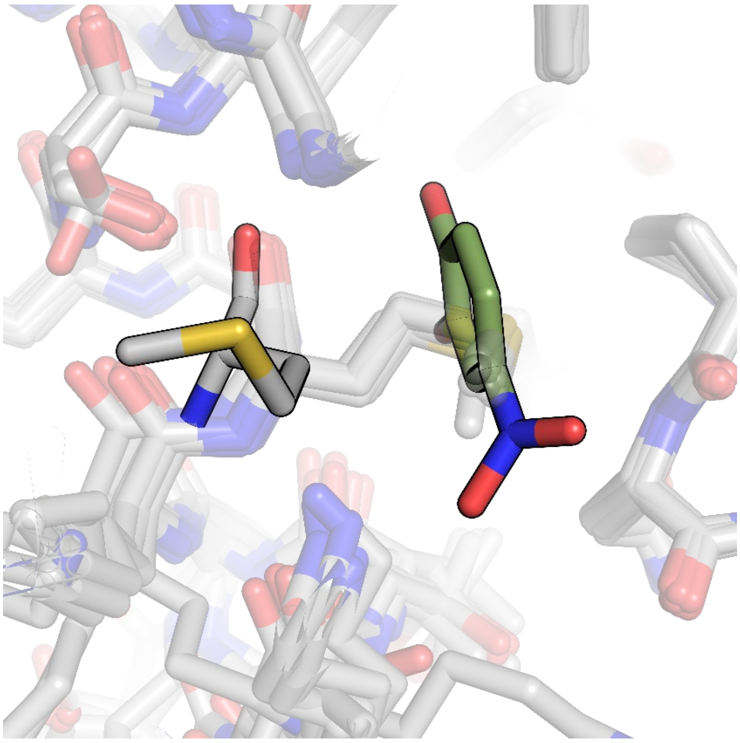

**Supplementary Figure 5: Displacement of Met278 by 26. 26** fragment is shown in green sticks. All fragment-bound crystal structures are aligned and shown translucent, with the reoriented Met278 from the **26**-bound structure outlined and opaque.

**Supplementary Table 6:** Crystallography statistics for heparanase bound to grown fragment C5A (**32**).

| **Structure ID** | C5A (**32**) |
|---|---|
| **PDB ID** | 9O2M |
| **Data collection** | |
| Space group | P $2_1$ $2_1$ $2_1$ |
| *a, b, c (Å)* | 59.31 74.97 124.65 |
| a, b, c (°) | 90 90 90 |
| Resolution (*Å)** | 47.93-2.00 (2.05-2.00) |
| $R_{merge}$ | 2.084 (27.439) |
| $R_{pim}$ | 0.881 (11.744) |
| $I/\sigma I$ | 4.80 (1.10) |
| $CC_{1/2}$ | 0.488 (0.486) |
| Completeness (%) | 98.7 (97.0) |
| Redundancy | 13.2 (13.2) |
| **Refinement** | |
| Resolution (*Å)* | 47.93-2.00 (2.05-2.00) |
| No. reflections | 38579 (2562) |
| $R_{work}$ | 0.1762 |
| $R_{free}$ | 0.2171 |
| No. atoms | 3984 |
| Protein | 3682 |
| Ligand/ion | 17 |
| Water | 285 |
| B-factors (overall) | 36.99 |
| Protein | 36.47 |
| Ligand/ion | 45.32 |
| Water | 43.24 |
| R.m.s. deviations | |
| Bond lengths (*Å)* | 0.005 |
| Bond angles (°*)* | 0.780 |

* Statistics for the highest resolution shell are shown in parentheses

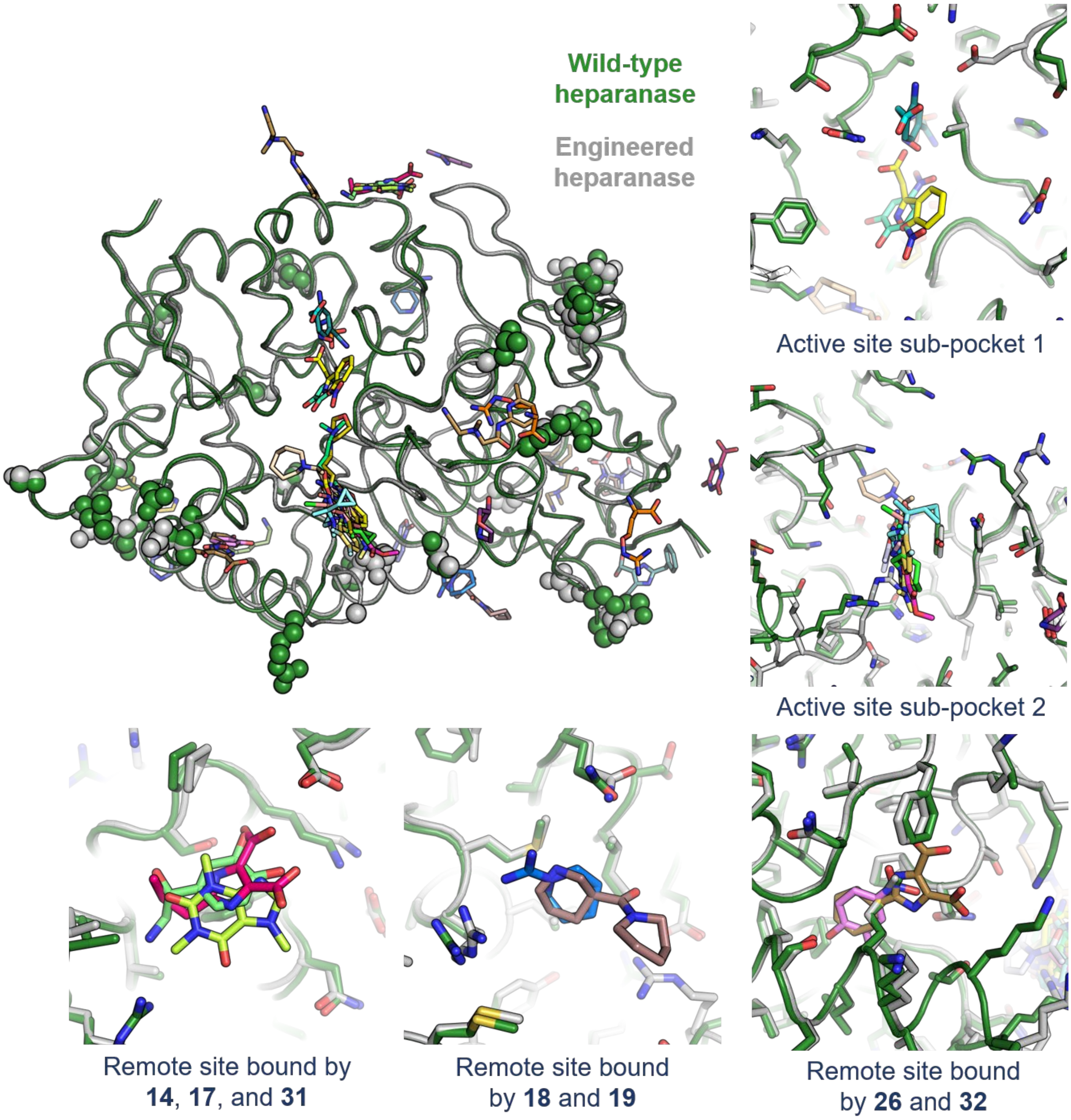

**Supplementary Figure 6: Fragment binding sites are distal to mutations introduced in the engineered heparanase variant.** Wild-type human heparanase (PDB: 5E8M) is shown in green, and engineered heparanase (PDB: 7RG8) is shown in grey. Residues mutated between the two sequences are highlighted in spheres, and fragments are shown in coloured sticks. Key binding sites are shown in individual panels.

**Experimental Procedures**

Protein expression and purification

An engineered heparanase variant designed previously[1] was used. The mutant heparanase P6 expression construct was co-transformed into *E. coli* Shuffle T7 Express cells (NEB), along with GroEL/ES + trigger factor chaperones in a pACYC vector and spread on an agar plate with ampicillin (100 mg/L) and chloramphenicol (34 mg/L). 1% overnight seed culture from a single colony was inoculated into 1 L of lysogeny broth (LB) medium supplemented with ampicillin (100 mg/L) and chloramphenicol (34 mg/L), before incubation at 37 °C for 5 hours until the optical density was between 0.8 and 1.2. Overexpression of heparanase was induced by adding 0.05 mM isopropyl β-D-1-thiogalactopyranoside (IPTG) and the culture was further incubated for 3 hours at 37 °C. The cells were harvested by centrifugation. The cell pellets were resuspended in buffer A (40 mM 2-[4-(2-hydroxyethyl)piperasin-1-yl]ethanesulfonic acid (HEPES) pH 8, 300 mM NaCl, 5 mM β-mercaptoethanol, 10% glycerol, 20 mM imidazole) with Turbonuclease (Sigma) and lysed by sonication (Omni Sonic Ruptor 400 Ultrasonic homogenizer). The lysate was filtered (0.45 µm) and loaded onto a nickel-nitrilotriacetic acid (Ni-NTA) column (GE healthcare) and eluted with 100% buffer B (buffer A + 500 mM imidazole). The peak eluent was loaded onto heparin affinity column (GE healthcare) equilibrated in buffer C (20mM HEPES pH 7.4, 200 mM NaCl, 1 mM DTT, 10% glycerol) and eluted with 100% buffer D (buffer C + 1.5 M NaCl). The peak eluent was loaded onto a size exclusion column (HiLoad 26/600 Superdex 200 pg, GE healthcare) and eluted in a buffer E (20 mM sodium acetate pH 5.0, 200 mM NaCl, 10% glycerol, 1 mM DTT). The final concentration of heparanase from the gel filtration was measured by absorbance at 280 nm using NanoDrop One (Thermo).

Protein crystallisation and structure determination

Apo heparanase crystals were obtained using the hanging-drop vapor-diffusion method, at 18°C, where a crystallisation solution (25% polyethylene glycol (PEG) 4K, 0.2 M ammonium sulfate and 0.1 M sodium acetate, pH 5.0) was added to each well of a 24-well Linbro crystallisation plate. Drops consisted of 0.5 µL of 7.77 mg/mL protein and 0.5 uL of crystallisation solution. Large, rod-shaped crystals grew overnight at 18 °C.

A JBS:Frag Xtal Screen (Cat.# X-FS-101) was used for fragment screening. A sitting-drop fragment tray was created using the computationally-identified fragments by adding 0.5 µL of fragment dissolved in DMSO at a concentration of 100 mM to the well and allowing the DMSO to evaporate at approximately 40 °C overnight. 30 µL of crystallisation solution was pipetted into the reservoir of the fragment tray, and 1 µL into the well containing the dry fragment to give a final fragment concentration of 50 mM. Apo heparanase crystals were selected based on size and soaked in the fragment tray for 10 to 20 minutes. The soaked heparanase crystals were then fished from the fragment tray and flash cooled in liquid nitrogen prior to analysis on the MX1 and MX2 beamlines at the Australian Synchrotron.[2, 3]

Diffraction data were processed using XDS[4] and DIALS[5] before scaling in CCP4's Aimless.[6]The search model used to solve the mutant structure was created using PDB ID: 5E9C[7] in CCP4's Phaser.[8] The search model generated from this for the mutant heparanase construct was used for molecular replacement in Phaser for all further crystal structures. The structures underwent iterative rounds of rebuilding in Coot[9] and refinement in Phenix.refine[10] before examination by PanDDA.[11]

The ground-state data set used in PanDDA was made up of 32 apo heparanase crystal structures that were auto-refined using the DIMPLE pipeline in CCP4 before further refinement using Phenix.refine.[6, 10] Coordinates and cif files for each of the fragments were generated using Phenix's eLBOW.[12]

In some structures there was clear electron density for fragments but for others it was ambiguous. In order to systematically identify if there was density for the fragments that was unique from apo structures of heparanase, the Pan-Dataset Density Analysis (PanDDA) method was used.[11] The PanDDA protocol as described by Pearce et al. (2017) was followed with appropriate parameters changed to suit the dataset being analysed.[11] The input datasets each had an RMSD of less than 0.6 Å to the reference structure. The PanDDA.analyse function was used to generate the mean density map for the ground-state dataset and identify unmodelled density in the sample data set that differs significantly from the ground-state. Identified unmodelled areas of density where then visually examined in Coot[9], fragments were modelled in and merged with the protein if the density looked appropriate using the PanDDA.inspect function.[9] Interactions between the fragments and the protein were found through visual examination and the 'environment distances' tool in CCP4's Coot[9] and figures were generated using Pymol.[13]

Computational docking

Docking was performed with the OPLS3e force field, and was completed in in Schrödinger 2020.3 Maestro v12.5.139.[14, 15]

*SiteMap*

PDBs of several inhibitor bound (6ZDM,[16] 5L9Z,[7] 5E98,[17] 5E97[17] and 5E9C[7]) and apo (5E8M[17]) heparanase structures were downloaded from the Protein Data Bank[18] and prepared through the Protein Preparation Wizard[19] function in Schrödinger. Epik[20] was used to define the residue protonation states at pH 5 ± 2, crystallographic waters were removed and the energy of the system was minimized. Site maps for the active site and potential allosteric sites for each of the protein structures were found using the SiteMap[21] feature in Schrödinger, with sites requiring at least 15 points per site, a restrictive definition of hydrophobicity and using a standard grid. Five potential allosteric sites were selected based on how many of the six protein structures they appeared in, the DScore, and the SiteScore. Receptor grids for each of these sites were prepared using Glide.[22] The default options for the scaling factor (1.0 Å) of the van der Waals radii of receptor atoms and the partial charge cut off (0.25) were used. Advanced settings were not changed. Figures of the site maps were generated using Pymol (v. 2.5.0).[13]

Identification of potential binding sites on the surface of heparanase were carried out using the SiteMap[21] package within Schrödinger with several inhibitor bound (6ZDM,[16] 5L9Z,[7] 5E98,[17] 5E97[17] and 5E9C[7]) and apo (5E8M[17]) heparanase structures. The main two metrics examined to identify likely binding sites were the SiteScore and the DScore of each site. Potential binding sites within a protein are identified by their size, exposure to solvent, and the hydrophobic and hydrophilic character of the site. The SiteScore provides a metric that takes these factors into account that can be used to easily compare the sites.[23] Similarly, the "druggability" of the identified sites is indicated by the DScore, where a higher DScore indicates a more easily druggable site. Like SiteScore, this score also takes into account the size of the site, its exposure to the solvent and the hydrophobicity/hydrophilicity of the site, but weighs them differently in order to produce a score that provides information into how easily the site can be drugged.[23] In order to be a promising druggable site for small molecules, the sites must have a low hydrophilicity, as well as a reasonable size and enclosure. Therefore, sites with a high SiteScore (>0.8) and DScore (>0.7) are indicative of sites with the desired druggability characteristics, and are considered good leads.[23] Ten potential allosteric sites were identified on the surface of heparanase. Sites A, B, C, D, and E were chosen for docking studies based on their relatively high SiteScore and DScore as shown in Supplementary Figure 2. Sites that were rejected from further studies had low scores, low conservation across the sample proteins, or a combination of these two factors.

*Molecular Docking Studies*

The Fragmenta (http://www.fragmenta.com/) library containing 2099 fragments was downloaded from the Zinc15[24] website and prepared through LigPrep[25] in the Schrödinger suite to generate the protonation states of the ligands at a pH of 5. The Fragmenta library was docked to the active site and the selected potential allosteric sites using High Throughput Virtual Screening (HTVS) in Glide. This was performed in triplicate to test for the reproducibility of the docking. The ligand sampling was set to rigid and the option to add Epik penalties to the docking score was included. The default options for the scaling of the van der Waals radii of the ligand (0.80 Å) and the partial charge cut off (0.15) were also used. Core and constraints parameters were left as default. The output of the HTVS was organised based on the Glide G-score and the top 250 results for each site were analysed. From this, only the fragments that were seen to bind in all three of the HTVS runs moved on to the next phase. The selected fragments from the HTVS were then docked to each of the sites using Standard Precision (SP), with the same parameters as HTVS and the output set to write at most five poses per ligand. The output data were processed in the same way as the results for HTVS, taking the top 50 results for each site. The top hits for SP were then docked to each of the binding sites with Extra Precision docking (XP) and the results were processed in the same way. 18 commercially available fragments were selected (Supplementary Table 4) across all six sites for validation through crystallography and inhibition assays based on the docking score and Glide G-score from the XP docking. The fragments were purchased from Sigma Aldrich and AK Scientific.

Fragments were selected for crystallography based on their G-score (kcal/mol). This score is representative of a fragment's binding free energy, where a more negative score is indicative of a tighter binding fragment.[26] Schrödinger's Glide module guidelines suggest that a good G-score is −10 kcal/mol or lower for ligands.[26] However, as fragments generally have a lower binding affinity than larger molecules, a slightly less negative G-score of −7 kcal/mol can be accepted as a good score for binding.

Enzyme inhibition assay

In order to determine the inhibitory activity of the fragments, a colorimetric assay designed by Hammond *et al* was used.[27] 96 well plates were treated with 1% bovine serum albumin (BSA) dissolved in phosphate buffered saline (PBS) containing 0.05% Tween-20 (PBST), and incubated at 37 °C for 75 minutes. After incubation, the plates were washed three times with PBST, tapped dry, then air dried for 10 minutes. The plates were sealed and stored at 4 °C until used.

The assay mixture contained 75 µL of 50 mM Na acetate buffer (pH 5.0), 10 µL of 8 nM heparanase in 0.001% Tween-20, 50 mM sodium acetate buffer (pH 5.0) and 10 µL of 1 mM fondaparinux (Aspen). 5 µL of 100 mM fragment solution dissolved in dimethyl sulfoxide (DMSO) was added to each well and a 1 in 2 serial dilution of the fragment in DMSO was carried out across the row, to make a total assay volume of 100 µL. The positive control contained no inhibitor, and the negative control had no inhibitor and no enzyme, with the inhibitor replaced with 5 µL of DMSO in both to give a total volume of 100 µL. A standard curve of a control inhibitor, BT2162 dissolved in DMSO, was used starting at a concentration of 125 µM, and diluted 1 in 2 by serial dilutions. The plates were incubated at 37 °C for 16–20 hours. After the incubation period 100 µL of 1.69 mM WST-1 in 0.1 M NaOH was added to each well. WST-1 is a cell proliferation agent that turns deep blue upon binding to the reducing ends of the hydrolysed fondaparinux. The plate was then resealed and developed by incubation at 60 °C for one hour, prior to measuring the absorbance at 584 nm. All assays were completed in duplicate.

(13) *The PyMOL Molecular Graphics System*; Schrodinger, LLC: 2021.
